# A suite of enhancer AAVs and transgenic mouse lines for genetic access to cortical cell types

**DOI:** 10.1101/2024.06.10.597244

**Authors:** Yoav Ben-Simon, Marcus Hooper, Sujatha Narayan, Tanya Daigle, Deepanjali Dwivedi, Sharon W. Way, Aaron Oster, David A. Stafford, John K. Mich, Michael J. Taormina, Refugio A. Martinez, Ximena Opitz-Araya, Jada R. Roth, Shona Allen, Angela Ayala, Trygve E. Bakken, Tyler Barcelli, Stuard Barta, Jacqueline Bendrick, Darren Bertagnolli, Jessica Bowlus, Gabriella Boyer, Krissy Brouner, Brittny Casian, Tamara Casper, Anish B. Chakka, Rushil Chakrabarty, Rebecca K. Chance, Sakshi Chavan, Maxwell Departee, Nicholas Donadio, Nadezhda Dotson, Tom Egdorf, Mariano Gabitto, Jazmin Garcia, Amanda Gary, Molly Gasperini, Jeff Goldy, Bryan B. Gore, Lucas Graybuck, Noah Greisman, Francoise Haeseleer, Carliana Halterman, Olivia Helback, Dirk Hockemeyer, Cindy Huang, Sydney Huff, Avery Hunker, Nelson Johansen, Zoe Juneau, Brian Kalmbach, Shannon Khem, Emily Kussick, Rana Kutsal, Rachael Larsen, Changkyu Lee, Angus Y. Lee, Madison Leibly, Garreck H. Lenz, Elizabeth Liang, Nicholas Lusk, Jocelin Malone, Tyler Mollenkopf, Elyse Morin, Dakota Newman, Lydia Ng, Kiet Ngo, Victoria Omstead, Alana Oyama, Trangthanh Pham, Christina A. Pom, Lydia Potekhina, Shea Ransford, Dean Rette, Christine Rimorin, Dana Rocha, Augustin Ruiz, Raymond E.A. Sanchez, Adriana Sedeno-Cortes, Joshua P. Sevigny, Nadiya Shapovalova, Lyudmila Shulga, Ana R. Sigler, La’ Akea Siverts, Saroja Somasundaram, Kaiya Stewart, Eric Szelenyi, Michael Tieu, Cameron Trader, Cindy T.J. van Velthoven, Miranda Walker, Natalie Weed, Morgan Wirthlin, Toren Wood, Brooke Wynalda, Zizhen Yao, Thomas Zhou, Jeanelle Ariza, Nick Dee, Melissa Reding, Kara Ronellenfitch, Shoaib Mufti, Susan M. Sunkin, Kimberly A. Smith, Luke Esposito, Jack Waters, Bargavi Thyagarajan, Shenqin Yao, Ed S. Lein, Hongkui Zeng, Boaz P. Levi, John Ngai, Jonathan Ting, Bosiljka Tasic

**Affiliations:** Allen Institute for Brain Science, Seattle, WA 98109; University of California, Berkeley, Berkeley, CA 94720; National Institute of Neurological Disorders and Stroke, National Institutes of Health, Bethesda, MD 20892

**Keywords:** Epigenetics, Transcriptomics, Enhancer, Cell type, Neocortex, AAV, Transgenic, Recombinase, Mouse, Human

## Abstract

The mammalian cortex is comprised of cells classified into types according to shared properties. Defining the contribution of each cell type to the processes guided by the cortex is essential for understanding its function in health and disease. We used transcriptomic and epigenomic cortical cell type taxonomies from mouse and human to define marker genes and putative enhancers and created a large toolkit of transgenic lines and enhancer AAVs for selective targeting of cortical cell populations. We report evaluation of fifteen new transgenic driver lines, two new reporter lines, and >800 different enhancer AAVs covering most subclasses of cortical cells. The tools reported here as well as the scaled process of tool creation and modification enable diverse experimental strategies towards understanding mammalian cortex and brain function.

## Introduction

To understand how the brain works, we need to establish the structure-function relationships between the units that comprise it and the outputs of its activity^1^. One approach is to gain experimental access to each building block (e.g., a cell type) and manipulate it to observe effects on an animal’s physiology and behavior^2–4^. Historically, the definition of building blocks and ‘tags’ for those building blocks have relied on non-systematic cell-type definition and non-systematic marker gene or genomic enhancer discovery. Numerous valuable tools have been developed and utilized to understand cell type functions in the brain, though they may not always achieve the desired level of cell-type resolution^2,3,5,6^.

Advances in next generation nucleic acid sequencing and machine learning have transformed cell classification based on single-cell genomics (transcriptomics, epigenomics, etc.)^7–16^. These new methods enable measurements of tens of thousands of molecular properties in individual cells and their subsequent classification into molecularly defined cell types. Cell type taxonomies have hierarchical organization, with cell type taxons of increased granularity frequently referred to as classes, subclasses, supertypes, and types/clusters^9,16^. Single-cell genomics also enables <u>cell type-congruent discovery</u> of marker genes and putative enhancers at all levels of the taxonomy^8,9,11,13,17–21^. These marker genes and enhancers can be used to make tools for precision cell type access, that can be employed to test the correspondence of molecularly defined cell types to ‘functional’ cell types^3,22–24^ (**Figure 1A**).

**Figure 1:**
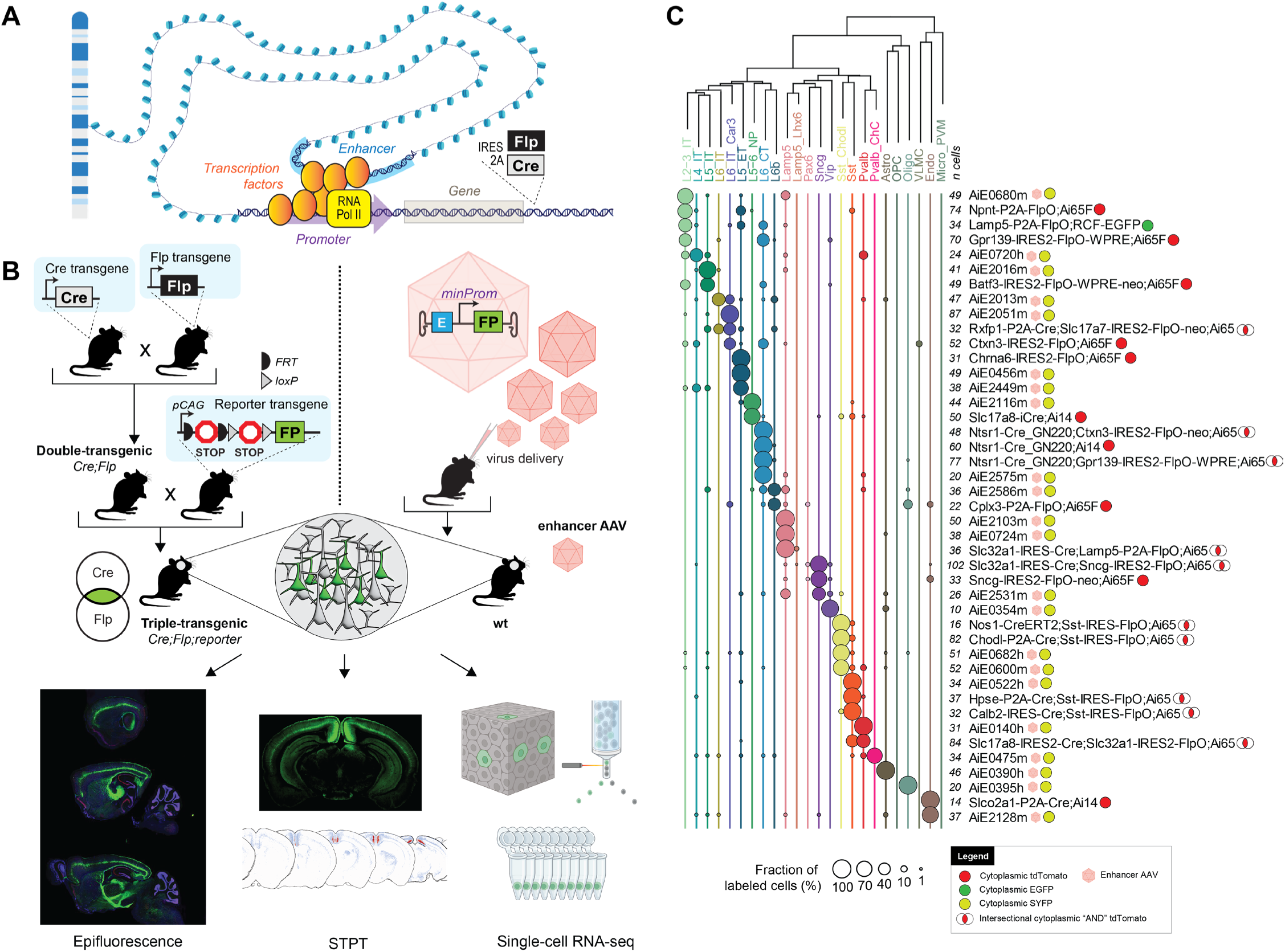
Cell-type-specific genetic tool design and characterization. **A.** A schematic illustration of gene regulation and location for recombinase insertions for knock-in transgenic mouse generation. **B.** Diagrams describing two strategies for genetic tool generation and characterization. *Left*: knock-in transgenic mouse lines are generated by insertions of recombinases into cell-type-specific differentially expressed genes. Generation of experimental animals requires one or more crosses to other recombinase lines and reporters. *Bottom*: The characterization of tool expression in brains of experimental animals is performed by three modalities: epifluorescence on select brain sections, serial-two-photon tomography (STPT) on whole brain, and single-cell RNA-seq on the visual cortex. *Right*: viral vectors utilize enhancers to achieve tool specificity. Generation of experimental animals requires retroorbital, intracerebroventricular or stereotaxic virus delivery to the animal. **C.** Single-cell RNA-seq data for some of the best tools reported in the study. Cortical taxonomy at the subclass level is on top and fraction of cells labeled per tool is represented by circles. Total number of cells per experiment (***n cells***) is represented in front of each tool’s name. Viruses are represented by pink hexagons. Other tools are transgenes.

Here, we relied on historical genomics data^17,18,25–27^, and newly-generated single-nucleus multiome data (snMultiome, 10x Genomics, consisting of joint snRNA-seq and snATAC-seq from each nucleus) to create a transcriptomic and epigenomic cross-species (mouse/human) taxonomy consistent with previously published datasets^17,18,25–27^. The newly generated snMultiome data were generated from the adult mouse cortex (somatosensory, motor, and visual areas), whereas the human cortical dataset was derived from the adult middle temporal gyrus^28^. We employed cell type definition in both transcriptomic and epigenomic space to discover corresponding ‘tags’: marker genes and putative enhancers to build and characterize a large suite of tools for cortical cell types.

We examined a total of 682 putative enhancers sequences: 599 novel and 83 that have been previously reported^17–19,29–31^. We tested these in mice by systemic introduction (retro-orbital injection) of individual enhancer AAVs expressing a fluorescent protein, employing the blood-brain-barrier penetrating PHP.eB capsid^32–34^. For initial evaluation of the enhancer AAVs, we visually examined and scored epifluorescence images of brain sections. Promising vectors were further evaluated for their brain-wide labeling pattern with serial two-photon tomography (STPT), and the cell-type specificity was determined by single-cell RNA sequencing (scRNA-seq) of labeled visual cortex cells (**Figure 1B**). We also attempted to optimize expression from these vectors by enhancer core bashing and concatenation (142 vectors), or expression of recombinase versions that can be combined with reporter mice (39 vectors)^17–19,35,36^. In total, we examined 863 enhancer AAVs in this study. Lastly, we generated new transgenic driver lines which can be used alone or in combination with enhancer AAVs to genetically access specific cortical populations at the finest taxonomical categories (supertype/cluster-level; **Figure 1C**).

In summary, we created and characterized a suite of genetic tools and compared them to select existing ones using a common battery of imaging and molecular techniques to provide detailed examination of their specificity and recommendations for their utilization (**Figure 1**). We are in the process of providing the tools through public repositories (Addgene and Jackson Labs) as well as adding data and metadata for all reagents (**Tables S1-S5**) to a new Allen Institute public web resource, the Genetic Tools Atlas (RRID:SCR_025643; https://portal.brain-map.org/genetic-tools/genetic-tools-atlas). These freely available public resources will enable scientists to access the information on these tools and select the best tool for their use.

## Results

### Selection of putative enhancer sequences from mouse and human genomics data

We selected putative enhancers based on a variety of data and diverse criteria over a span of several years. To identify putative enhancers, we used various chromatin accessibility genomics data for mouse and human cortex, some published^17,18,25–27^, some newly created and analyzed (see below). Most putative enhancer selection was based on 1) high and differential chromatin accessibility across the cortical cell subclasses and 2) proximity to marker genes. Putative enhancer sequence conservation across species was not a generally used criterion.

To summarize all the chromatin accessibility data across cell types and represent them in a unified way, as well as select more putative enhancers, we generated a new cortical snMultiome (10x Genomics, bimodal single-nucleus RNA-seq and ATAC-seq) dataset of single nuclei isolated from adult mouse visual, somatosensory, and motor cortices (82,654 nuclei post-QC; **Figure 2A**). We also employed a recently-published snMultiome dataset of nuclei isolated from adult human middle temporal gyrus (MTG, 83,977 nuclei post-QC; **Figure 2A**, right)^28^. For mouse, we mapped the single-nucleus transcriptomes to the recently-published whole mouse brain taxonomy (AIT21) using hierarchical approximate nearest neighbor (HANN) mapping^16^. For the human MTG data, single-nucleus transcriptomes were mapped to the great apes taxonomy with scANVI^37,38^. We compared the great ape taxonomy and a previously published mouse cortical taxonomy^9^ to generate a common list of cell types that occur in both species. This list was used to re-annotate mapped mouse and human nuclei at an intermediate cell type resolution (cell subclass). This annotation was performed on the transcriptomic part of the multiome datasets and it resulted in comparable cell populations across mouse and human datasets; similar to what has been described before^37,39^.

**Figure 2:**
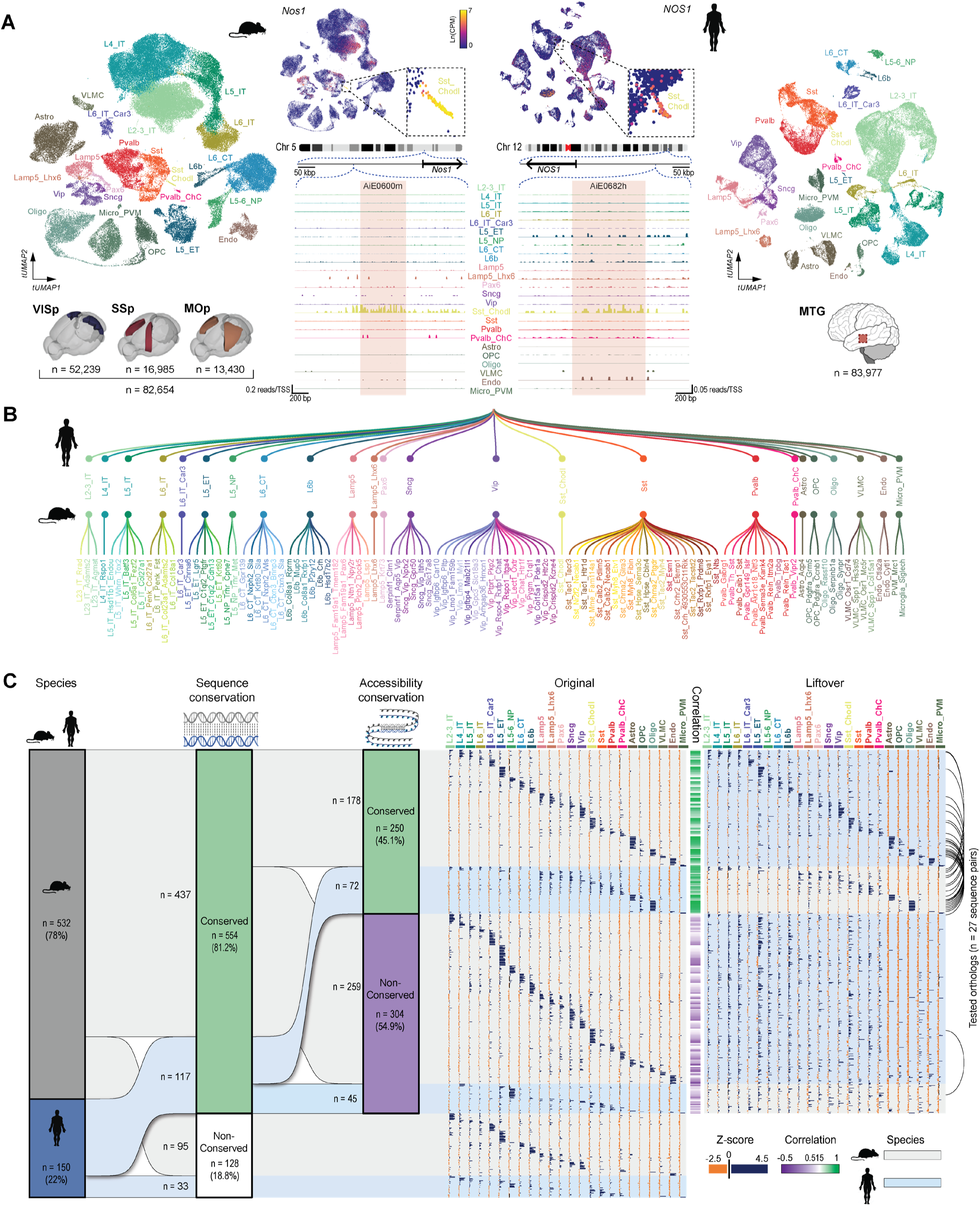
Selection of putative enhancer sequences. **A.** Mouse and human single-nucleus transcriptomes obtained from single-nucleus multiomes represented in a transcriptomic UMAP and labeled according to cortical cell subclasses. Mouse nuclei were collected from the primary visual (VISp), somatosensory (SSp) and motor (MOp) cortices; human nuclei were collected from the middle temporal gyrus (MTG). Numbers of nuclei included in the analysis are represented by ‘n’. **B.** Simplified representation of the unified mouse-human taxonomy of cortical cell subclasses, along with the cluster-level taxonomy for mouse only. **C.** Summary of all ‘native’ putative enhancer sequences tested (n = 682), divided by the genome of origin, followed by cross-species conservation of sequence and accessibility (left). The modified sequences produced by concatenation are not included. The ‘Original’ and ‘Liftover’ plots show relative accessibility of all individual sequences in each subclass in their respective species, alongside the relative accessibility of its orthologous liftover sequence in the other species, respectively. Correlation between the original and liftover accessibility data is shown as a green/purple heatmap in the middle of these plots. Black arcs on the very right indicate instances where orthologs from both species were tested.

The transcriptome-based annotation for each nucleus was used to label its respective ATAC-seq profile for downstream analysis. Pseudobulk ATAC-seq data were generated with ArchR^40^ and used to identify putative enhancers as differentially-accessible peaks across subclasses by MACS2 and ArchR^40,41^. As an example, we show gene expression data for the *Nos1* gene and the accessibility of two orthologous enhancers targeting Sst_Chodl cell type near this gene in the mouse and human datasets (**Figures 2A, 2B**).

In total, we selected 532 sequences from the mouse genome (average length ± SD = 583 ± 256 bp; range = 133-2096 bp, **Figure S1A**) and 150 sequences from the human genome (average length ± SD = 484 ± 176 bp; range = 162-1070 bp; **Figures 2C**, **S1A**). Each putative enhancer sequence was assigned a unique ID, consisting of the initials AiE for enhancers discovered at the Allen Institute, or ExE for enhancers previously reported by other groups^29,30^, followed by a four-digit number and a species indicator (“m” or “h” for mouse or human, respectively). To provide consistency to the enhancer collection, we applied the same naming structure to the previously published enhancers from Allen Institute^17–19^ and provide the original names as aliases (**Table S4**). Additionally, each enhancer was associated with one or two target cell populations (TCP), which allow us to predict which population/s they are most likely to be expressed in. These TCPs were determined by scaling ATAC signal across subclasses and nominating the subclasses with accessibility Z-scores higher than 2 as a TCP.

We also evaluated the degree of cross-species sequence conservation using the UCSC genome browser LiftOver tool^42^, followed by BLAST alignment, and correlated the chromatin accessibility pattern of the orthologous pairs, across the cortical subclasses (**Figure 2C**). Most of the selected candidate enhancer sequences were conserved between the two species (n = 554, 81.2%, **Figures 2C, S1A-B**). Among this conserved subset, 59.2% (n =178) of mouse and 39.0% (n = 72) of human enhancers also displayed conserved chromatin accessibility across species (**Figure 2C**). This dataset also included 27 tested orthologous pairs from the two species (black arcs in **Figure 2C**).

### Enhancer AAV screening *in vivo*

Candidate enhancer sequences were amplified from the respective genome and cloned into the previously-described AAV plasmid backbone^17,18^, upstream of a minimal promoter (beta-globin, rho or CMV), driving expression of the yellow fluorescent protein, SYFP2^43^. Most plasmids contained the beta-globin minimal promoter (MinBG; 94.6%). The plasmids were packaged into blood-brain-barrier-penetrating, PHP.eB-pseudotyped, AAV vectors^33,34^ and delivered retro-orbitally (RO) to ∼30 day-old wild-type mice of C57BL/6J background (average age ± SD = 30.8 ± 5.0 days, range = 27-70 days; **Methods**)^44^. About four weeks after virus delivery (average duration ± SD = 27.2 ± 1.5 days, range = 23-39 days; **Methods**), the brains were extracted, fixed, and sectioned along the sagittal plane, with five selected sections spanning the mediolateral axis mounted and imaged by an epifluorescence microscope (**Figure 3A**). We evaluated each experiment by visual inspection and categorized labeled cell populations (LCPs) in the cortex into 11 visually distinguishable categories (**Figure 3B**). Each LCP was further visually assessed for labeling density relative to the expected density for that target cell population (high vs. low density), as well as relative labeling brightness (high vs. low brightness; **Figures 3B, S2A**, **S2B**). Although we scored each LCP for both parameters, we considered brightness to be a more relevant feature of the enhancer than the density, as this factor is more easily comparable across different populations of labeled cells, and is more routinely used to evaluate enhancer strength^45–47^. In addition to annotating the neocortical populations, we also scored all labeled regions, brain-wide, using Allen Mouse Brain Reference Atlas naming scheme^48^ as these enhancer AAVs could be useful tools for other brain regions. For brevity, we are reporting only the neocortical labeling patterns here. The Allen Genetic Tools Atlas web portal (in preparation) will contain labeling scores for all tools and all brain regions.

**Figure 3:**
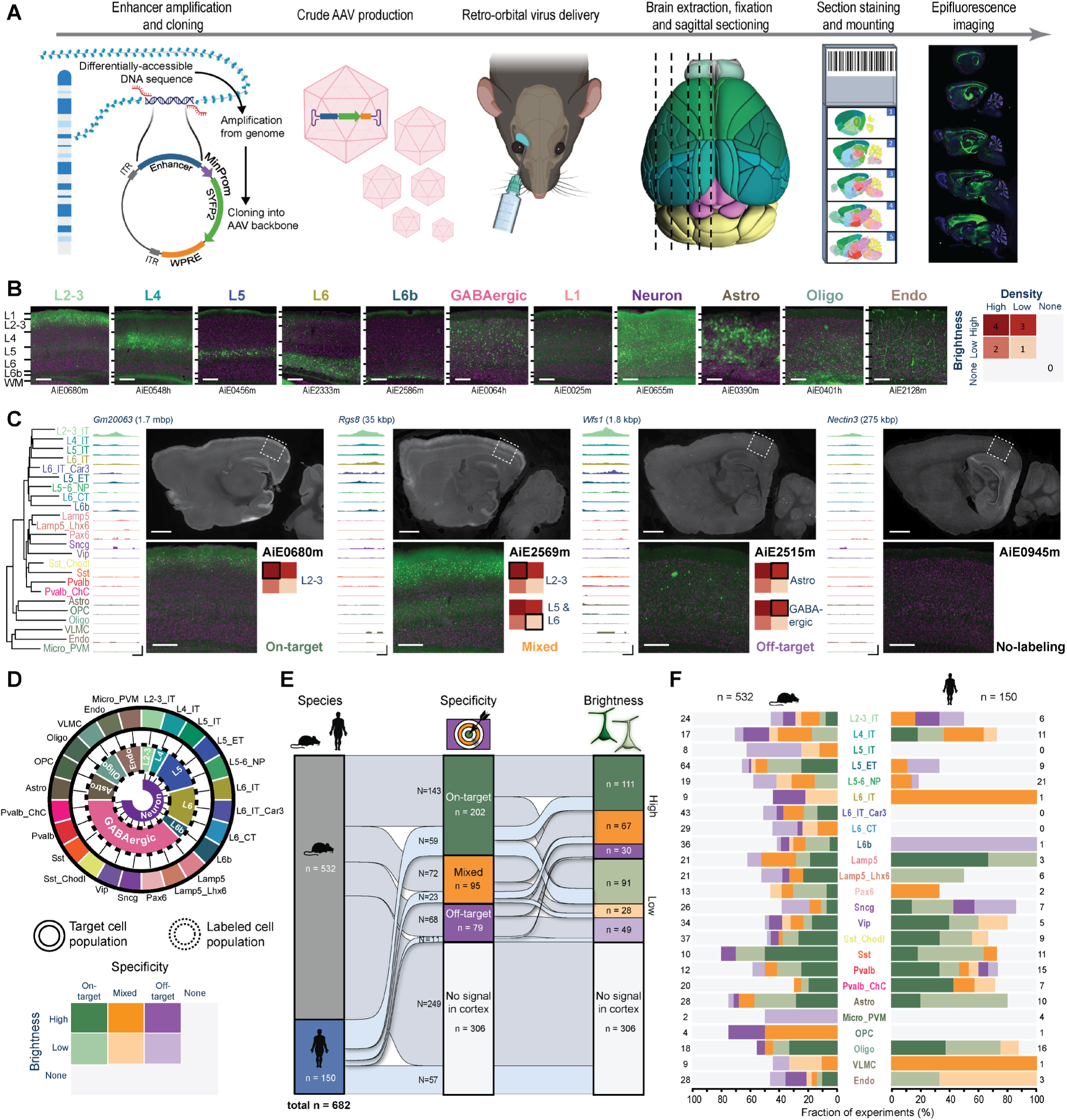
A pipeline for enhancer AAV screening in the mouse brain. **A.** Graphical summary of the enhancer screening pipeline. **B.** Representative images of the visual cortex, showing examples of 11 categories of visually distinguishable cortical populations used to evaluate the labeling pattern of each enhancer AAV. Individual enhancer IDs are shown beneath each image (left), and a scoring matrix for evaluating brightness of the fluorescent signal and the labeling density, compared to the density expected for each category (right). **C.** Representative tracks of chromatin accessibility for four individual enhancers targeting across the cortical subclasses, demonstrating differential accessibility in L2-3_IT, alongside representative epifluorescence image sets, showing the resulting labeling pattern, and the score given to each in VISp. The closest L2-3_IT marker gene is shown above each set of tracks, along with the distance from the enhancer to its TSS. Scale bars below the tracks represent 100 bp (horizontal) and 0.3 RPKM/cell (vertical). **D.** Schematic describing the approach used to determine target specificity, according to the alignment between the TCP and LCP (top left) and a matrix used for classifying all enhancers, based on a combination of their target specificity and signal brightness (bottom left). **E.** Summary plot showing performance of all tested enhancers (n = 682) according to the categories (right). > 50% of enhancers (n = 376) exhibited signal in the cortex, ∼43% were putatively on-target or had mixed (on- and off-target) labeling, and ∼30% were putatively on-target (202/682). **F.** Proportion of enhancers in each of the categories specified in (D), according to their genome of origin (top, n = 150 for human and 532 for mouse) and TCP. Numerical values represent the number of tested enhancers in each column. For images, scale bars for full section and expanded views are 1.0 and 0.2 mm, respectively.

Since cell type distribution and cell shapes across neocortical layers for all TCPs are known^9,49,50^, we visually matched each observed LCP with the expected pattern of the TCP to derive initial evaluation on target specificity. Following this TCP-LCP matching, we assigned each enhancer to 1 of 7 categories, indicating whether it had putative on-target, mixed, or off-target labeling, or whether no signal was detected; each of the first three categories was further divided according to the labeling brightness (**Figure 3C, 3D**). Multiple TCPs can belong to a single LCP, for example, in the case of different subclasses of GABAergic neurons. Therefore, this level of analysis can only provide putative on-target specificity for groups clearly differentiable by visual inspection. However, as presented below, this scoring scheme identified many highly specific enhancers and provided a nomination path for their additional characterization. Of the 682 unique enhancers screened, ∼55% produced labeling in the cortex, ∼43% were putatively on-target or had mixed (on- and off-target) labeling, and ∼30% were putatively on-target (**Figure 3E**).

We also used the scoring data to examine the distributions of scores across cell subclasses and species, as well as the degree of sequence/accessibility conservation (**Figure 3F, S3A** – **S3D**). For human enhancers, we detect a statistically significant lower-on target rate for glutamatergic cell class (**Figure S3A**, top). We observe a similar trend for mouse enhancers, but it did not reach statistical significance (**Figure S3A**, top). We also observe a higher on-target rate for enhancers with conserved sequence and accessibility (**Figure S3A**, bottom). Specifically, we note that the precise degree of conservation, measured by the fraction of overlapping base pairs (**Figure S1E**) strongly affects both the specificity and brightness of the enhancer (**Figure S3B**). When comparing orthologous pairs, no significant differences in score distribution were observed overall (**Figures S3C**). However, high variability within the categories was noted, showing that the differences in orthologous sequences can affect enhancer specificity and strength (**Figure 3D**). We also found that orientation of enhancers in the context of AAV appears to have no effect on their performance, (**Figures S3E, S3F**), as has previously been observed for the SV40 enhancer^51^.

In summary, we describe a screening pipeline for putative enhancer sequences in the mouse brain and conclude that sequences derived from mouse and human genome can perform comparably well at the subclass cell type level, when evaluated in the mouse brain, with roughly 30% of them showing on-target specificity as determined by our manual scoring.

### Enhancer AAV secondary characterization

Enhancers that showed promising results in our initial screen were nominated for further evaluation with single-cell RNA-seq (scRNA-seq) with Smart-seq v4 (SSv4) and/or serial two-photon tomography (STPT)^52^ (**Figure 4A**). To better evaluate target specificity and labeling brightness, we delivered each vector with the same parameters used for the primary screen, and 4-8 weeks later (average duration ± SD = 45 ± 10 days; range = 21-91 days), dissected the visual cortex, dissociated the cells and isolated SYFP2(+)/DAPI(-) cells with FACS (**Figure S4A**). These cells were subjected to scRNA-seq and their transcriptomes were mapped to the mouse VISp cortical taxonomy (**Methods**). These data enabled us to quantify the fraction of cells belonging to any cell type for each experiment (**Figures 4B, 4C**, **S4B**, **S4C**). In addition, we quantified the expression level of the SYFP mRNA (median SYFP2 mRNA count per million transcripts) and normalized it for each cell type to the median expression of *hSyn1*-promoter/enhancer-driven SYFP2 mRNA (**Figure 4B, S4D**).

**Figure 4:**
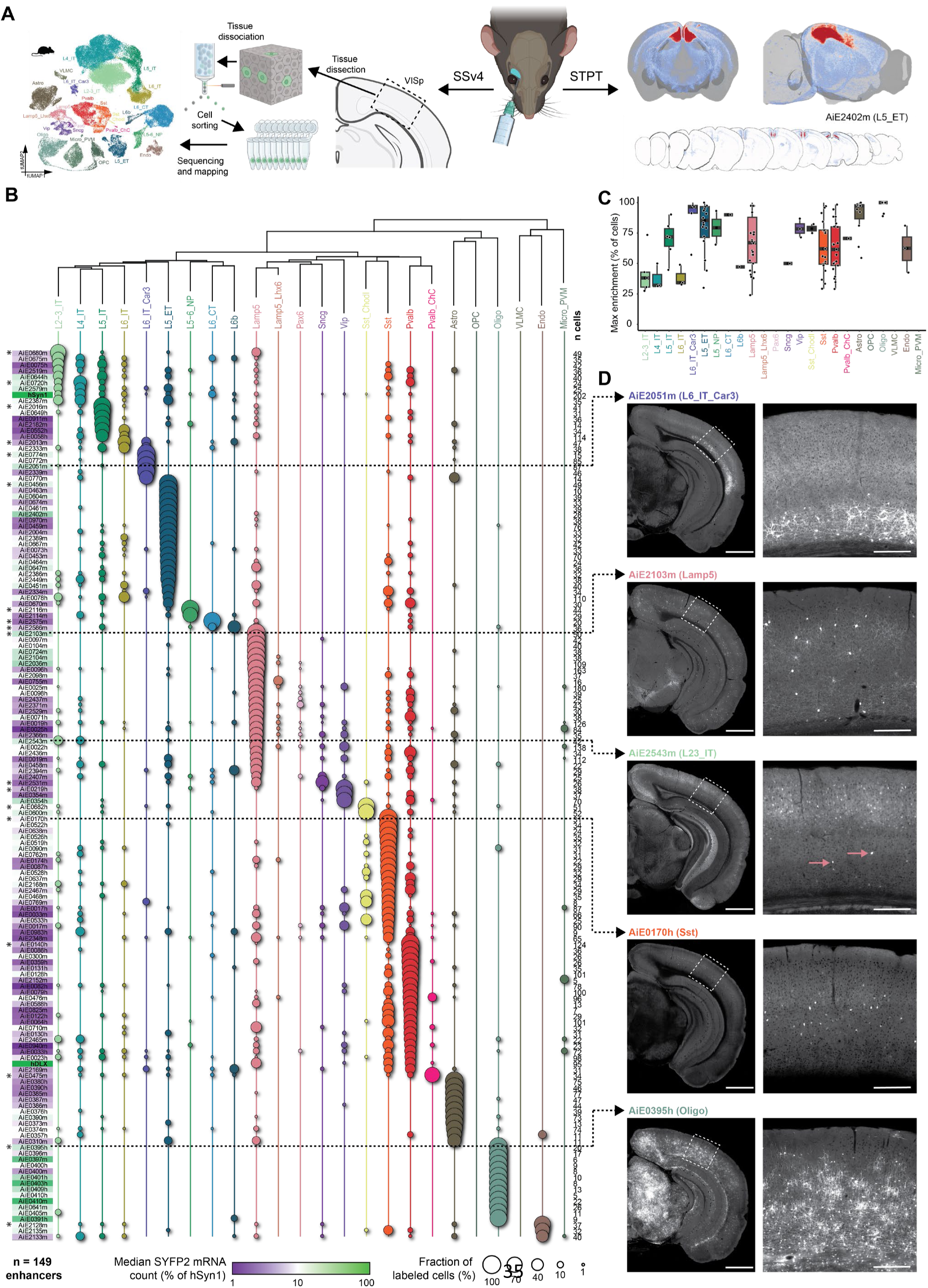
Secondary validation of target specificity, with scRNA-seq and whole-brain imaging. **A.** Schematic of methods for secondary validation. **B.** scRNA-seq analysis using SSv4, of FACS-sorted SYFP2^+^ cells from the mouse visual cortex, following RO administration of the enhancer AAV. The fraction of cells mapped to each cortical subclass corresponds to circle size, and the median SYFP2 mRNA count in each experiment, relative to the hSyn1 promoter, is denoted by a purple-to-green color gradient. The number of sequenced cells for each experiment is shown to the right of the table. Total number of enhancers examined is n = 149. Asterisks denote the top performing enhancers for each subclass, i.e., the ones with highest proportion of cells mapping to the subclass of interest. **C.** Box plot showing for each cortical population, all enhancer AAVs for which that population was the main enriched target population. The thick black bars represent medians, color-coded boxes represent top and bottom 25%, and whiskers represent top and bottom 10%. Data for individual enhancers is shown as superimposed black circles. **D.** Representative STPT images for five enhancers, with an expanded view of VISp displayed to the right. Dashed arrows connect each image set to its corresponding SSv4 data. In the case of AiE2543m, which labels L2-3_IT cells and Lamp5 cells, pink arrows point to sparse, yet brightly labeled non-L2-3 neurons, which are likely the Lamp5 interneurons. These are overrepresented in SSv4 (B), likely due to the stringent gating strategy in FACS focusing on the highly fluorescent cells. For images, scale bars for the hemisphere and the VISp magnified view are 1.0 and 0.2 mm, respectively.

We find that our initial screen produced many highly specific enhancer AAVs: more than 50% of the enhancers examined with scRNA-seq exhibited target specificity of >70% at the subclass cell-type level and 28% were >90% specific (**Figure S4C**). However, we note that SYFP2 mRNA expression driven by the enhancer AAVs was relatively low for most of the enhancers, when compared with the pan-neuronal *hSyn1*-promoter/enhancer (average ratio ± SD = 13.3 ± 15.3% of hSyn1; **Figure S4D**). We also observed that ∼29% of enhancers exhibited labeling distribution patterns that were not significantly correlated with the distribution of accessibility across subclasses (**Figure S4E**). This unexpected result could be partially explained by the way cells were collected and sorted for sequencing: To avoid false negatives in FACS, we set the sorting gates relatively stringently, to collect the brightest cells (**Figure S4A**)^19^. However, this can lead to biases in the results, in cases where two or more populations are labeled, with the more abundant population being dimmer than the smaller one, as in the case of AiE2543m (**Figure 4D**). Note that evaluation of specificity by scRNA-seq of FACS-isolated cells can also lead to biases in cases where labeled population includes cell types particularly sensitive to the dissociation and sorting process (see below)^8,9^.

For enhancers with putative on-target labeling patterns, we generated whole-brain STPT image sets to further interrogate the enhancer AAV labeling patterns. Compared to five epifluorescence images, STPT enables more complete evaluation of the entire cortex. STPT also examines all other parts of the brain and shows if the enhancer is cortex-specific or active in other brain regions. Finally, brain-wide visualization reveals axonal projection patterns and can help distinguish between different types of cortical projection neurons. We examined ∼34% (180/532) of mouse enhancers and ∼41% (62/150) human enhancers for a total of 242 STPT whole-brain image datasets (representative examples in **Figure 4D**). These data complement the SSv4 data (**Figure 4B**), as well as confirm and extend primary screen data, as they cover the whole brain with a series of coronal images. In all cases, we find that the STPT data are in full agreement with the labeling patterns observed in the primary screen.

We cross-correlated the specificity of each enhancer, measured as the maximal fraction of labeled cells, with the SYFP2 transcript count and cross-species conservation of sequence and accessibility (**Figure S4F**). We find that enhancer specificity was positively correlated with the SYFP2 transcript count, suggesting that specific enhancers also tend to drive stronger cargo expression. We also found a significant positive correlation with the mouse-human sequence homology for the 130 enhancers for which the sequence was conserved, but not with the degree of accessibility conservation across the two species (**Figure S4F**). It should be noted that this analysis was performed only on the SSv4 data, which comprise only enhancers with demonstrated putative on-target specificity in our primary screen, and therefore is not representative of the entire collection.

These analyses highlighted enhancers which delivered high specificity as potentially useful tools for selective targeting of cortical cell types. The relevant experimental data for SYFP2-expressing vectors in this study were organized in supplementary material (**Documents S1-S5**) and will be available at the Genetic Tools Atlas web portal (RRID:SCR_025643; https://portal.brain-map.org/genetic-tools/genetic-tools-atlas).

### Enhancer optimization and use diversification

Previous studies have shown that the active site of a putative enhancer sequence can be short and can be discovered through an approach called “bashing”, where shorter fragments of the originally tested sequence are individually examined for their ability to drive cargo expression^53–56^. We proceeded to fragment a subset of enhancers into three putative “cores” (C1, C2, and C3; average length ± SD = 212 ± 75 bp) that tile the originally-examined sequence with 50 bp overlap^17^. To try to enhance labeling brightness and specificity, we cloned a 3x-concatenated version of each core (3xCore, 3xC) into the AAV backbone (**Figure 5A**). We have previously shown that enhancer concatenation can increase reporter expression^17–19^. In addition, this approach may also increase labeling specificity in cases where the complete enhancer sequence contains several independent elements, each driving expression in a different population of cells^57,58^.

**Figure 5:**
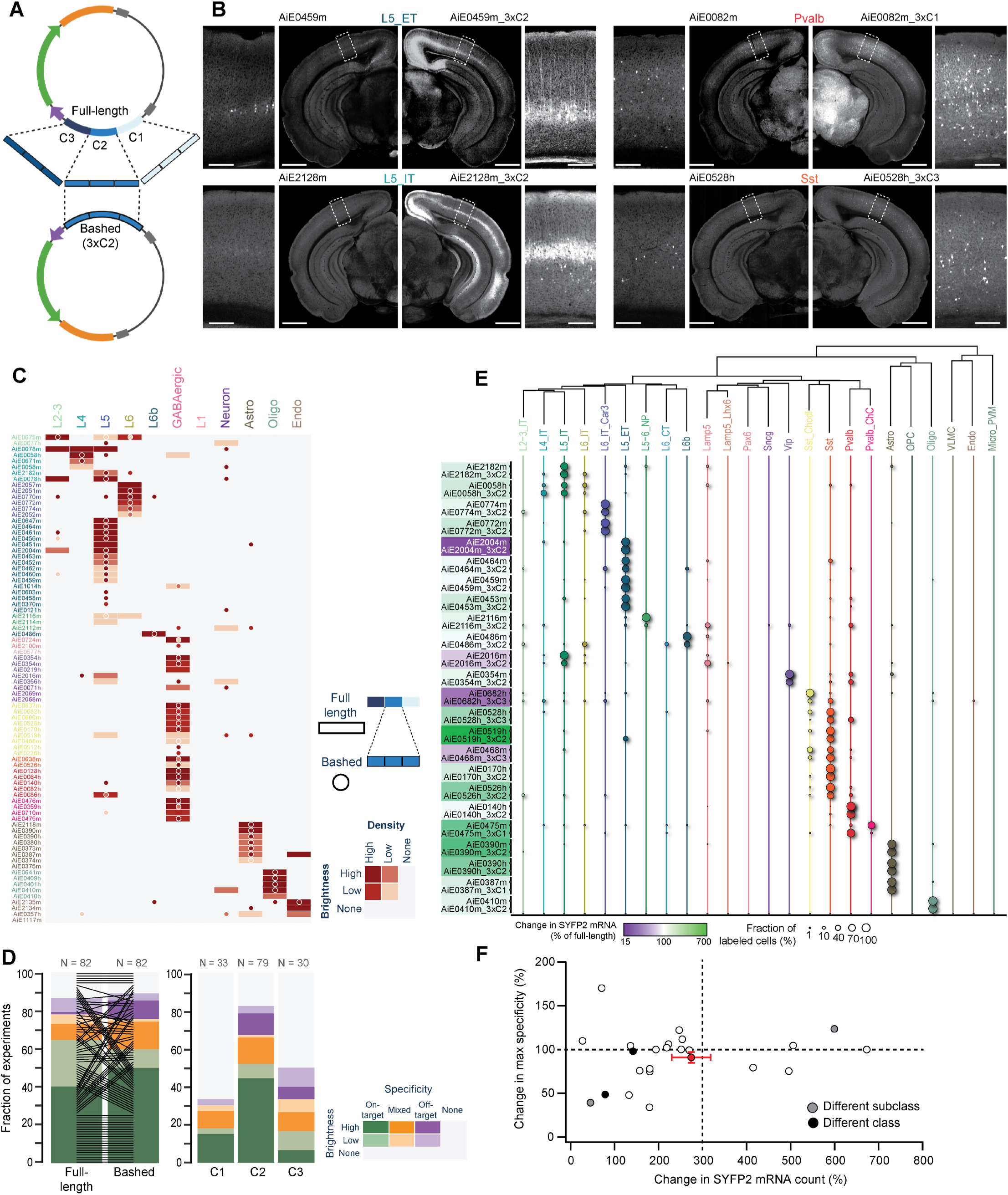
Optimization of enhancer activity through core bashing. **A.** Schematic representation of the core bashing approach for enhancer optimization (C = Core). **B.** Representative STPT images of coronal hemisections, showing labeling pattern for four individual full-length enhancers (left) and their best bashed version (right), which was selected according to the combination of brightness and specificity. A magnified view of VISp is shown alongside each hemisection. Scale bars for full section and expanded view are 1.0 and 0.2 mm, respectively. **C.** Heatmap showing the scoring results of epifluorescence image sets of the full-length enhancer (rectangles) alongside its best bashed version (circles); n = 82 pairs. **D.** Summary of the scoring data in (C), sorted according to brightness and specificity of the full-length vs. the bashed enhancer (left) and for the different cores tested (right). **E.** Dot plot of SSv4 data for full-length enhancers and their bashed counterpart shown in pairs, with circle size denoting the fraction of cells mapped to each of the cortical subclasses in each experiment. The color overlaying each pair name corresponds to the relative change in SYFP2 transcript count of the bashed relative to the full-length enhancer; n = 24 pairs. **F.** Change in specificity vs. change in SYFP2 transcript count for all enhancer pairs in (E). Average and SEM for all experiments corresponds to the red dot with error bars. Pairwise comparisons for individual enhancers correspond to white dots if no change in specificity if observed. If the bashed version preferentially labeled a different subclass or class compared to the corresponding full-length enhancer, the dots are grey or black, respectively.

We core-bashed a total of 82 enhancers, creating 3xC of all three cores or only the middle core (C2), which usually aligns with the peak of chromatin accessibility. For each original enhancer, at least one, and in some cases two bashed enhancers displayed labeling in the original TCP. Most also resulted in a marked increase in brightness (**Figure 5B**), particularly those core-bashed enhancers designed using the middle (C2) core. However, core-bashed versions occasionally labeled additional cell populations, which were not labeled by the original enhancer (**Figures 5C, 5D**). It is unclear if the concatenation of the cores produced novel TF syntax, which led to *de novo* labeling, or whether these populations were labeled by the full-length (original) enhancer but at levels below detection threshold. To further compare the select bashed versions with the original enhancers, we collected SSv4 scRNA-seq data from VISp. We compared the labeling specificity and SYFP2 mRNA expression levels of the concatenated bashed enhancer with the original enhancer. The behavior of individual enhancers varied, but on average, we observed ∼3-fold increase in SYFP2 mRNA count for the bashed 3x concatenated enhancer (average ratio ± SD = 274 ± 217% of full-length enhancer) with a slight reduction in the specificity (average ratio ± SD = 91 ± 30% of full-length enhancer; **Figures 5E, 5F**). This is in agreement with previous studies that show that enhancers can support target gene expression in additive fashion, as well as sub- and supra-additive fashion depending on the enhancers themselves^59,60^.

To diversify the use of our enhancer-based genetic toolkit, we replaced SYFP2 with a Cre recombinase, which would enable expression of a variety of tools from existing or new Cre-dependent AAVs or mouse transgenes. We employed iCre or a mutated iCre(R297T) with reduced recombination efficacy^61^ and removed the WPRE sequence in a subset of enhancer vectors. The latter two choices were implemented to counter loss of specificity that was more frequently observed in constructs with iCre and WPRE compared to the original constructs with fluorescent proteins, as reported previously^22^. We evaluated the recombination-mediated reporter labeling and compared it to the original pattern defined by constructs with SYFP2 (**Figure 6A**). Several enhancers labeling various TCPs showed highly similar labeling patterns following replacement of the fluorophore with a recombinase. Others produced broad labeling patterns (**Figures 6B** – **6D**) that could be overcome by using the mutated version of Cre, and/or by titrating the amount of virus delivered (**Figure 6E**).

**Figure 6:**
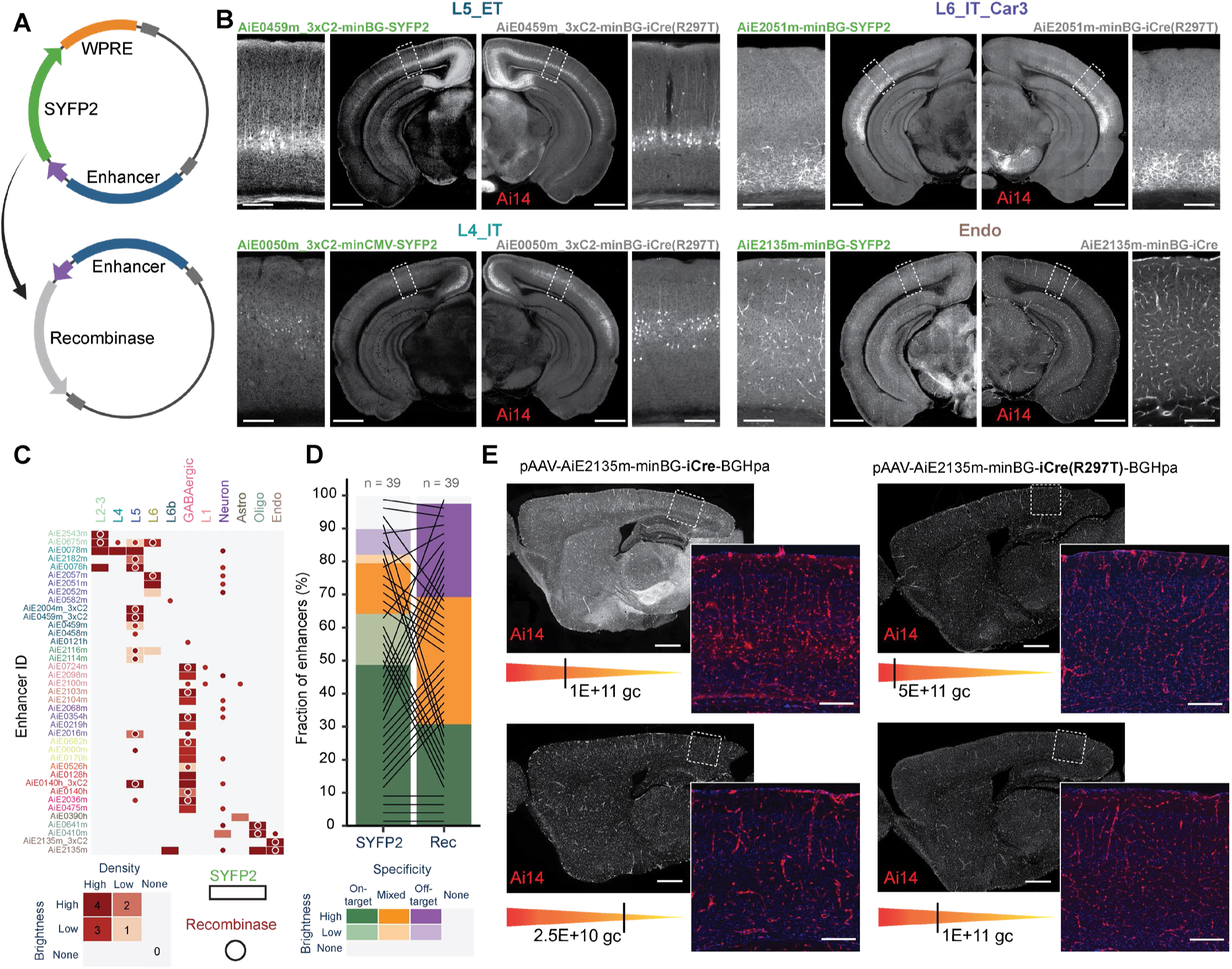
Recombinase-expressing enhancer-AAVs. **A.** Schematic representation of the vector design. **B.** Representative STPT images of coronal hemisections, showing labeling pattern for four individual enhancers expressing SYFP2 delivered to a wild-type mouse (left) and the same enhancers driving iCre(R297T), delivered to the *Ai14* reporter mouse (right). A magnified view of VISp is shown alongside each hemisection. **C.** Heatmap showing the scoring of epifluorescence imaging data of the enhancers driving SYFP2 (rectangles) or a recombinase (circles); n = 39 pairs. **D.** Summary plot of the scoring data in (C), comparing brightness and specificity of the SYFP or recombinase expression. **E.** Specificity of Cre-dependent recombination in endothelial cells with the Ai2135m enhancer is reduced in iCre compared to the mutated version iCre(R297T) and is more sensitive to the viral dose (gc, genome copies). Scale bars in (B) and (E), 1.0 and 0.2 mm for full section and expanded view, respectively.

To further test the applicability of these viral tools, we evaluated their performance under different delivery methods. We observed that intracerebroventricular (ICV) delivery of vectors into the lateral ventricle of newborn mouse pups can lead to specific and widespread labeling, comparable with RO injections, but with substantially brighter fluorescent signal^62^ (**Figures S5A – S5C**). However, in several cases, we observed a non-uniform signal distribution and an increase in non-specific labeling, which may have been present below detection levels when the virus was RO-delivered^62^. Interestingly, we found that some populations, particularly endothelial cells, could only be labeled when the virus was delivered RO, but not by ICV administration (**Figure S5C**), which is consistent with previous observations where alternative approaches were tested to target this population^21,33,63^. It has also been reported that certain cell subclasses such as VLMCs, OPCs and microglia have virtually absent transduction with PhP.eB-serotyped AAVs and hence are difficult to target via our current screening strategy, which exclusively uses the PhP.eB capsid for all vectors^64^. All the vectors designed for these cell subclasses led to off-target/mixed-target labeling (**Figure 3F**) and were not further investigated by SSv4.

In addition, we delivered several purified enhancer AAVs stereotaxically to the visual cortex and noted that the fluorescent signal was substantially brighter, restricted to the injection site, and mostly maintained specificity (**Figures S6A, S6B**). We also note that enhancers that exhibited weak fluorescence when delivered RO displayed increased signal intensity when delivered stereotaxically, likely due to higher multiplicity of infection (MOI) at the target region (**Figure S6A**). Finally, we provide a summary of ‘hall-of-fame’ enhancer tools with highest available strength and specificity of labeling for cell subclasses or clusters (**Table S5**).

### New transgenic lines targeting cortical cell types

Based on the transcriptomic taxonomies of cell types in the mouse cortex^8,9,13^, we selected marker genes that could label specific cell type taxons (subclasses, supertypes or clusters). We targeted regions of the taxonomy for which tools did not exist. Based on select marker genes, we generated a total of 15 new transgenic driver lines: 12 preferentially for glutamatergic and 3 preferentially for GABAergic cell types (**Figure S7A, 7, 8**). We also made two new reporter lines that address the unmet need for Cre “AND/OR” Flp reporters: *Ai193*^65^ and *Ai224*^66^ (**Figure S7B**).

Initial characterization for the new driver lines was performed by crossing to a fluorescent reporter line (*Ai14* for Cre, *Ai65F* for Flp transgenes, respectively)^67,68^ and subsequent STPT on the whole mouse brain. We examined labeling of cortical cells for expected cell distribution (**Table S1**). Most driver lines produced expected labeling based on cell types expressing the targeted marker gene and were further analyzed by single-cell transcriptomic profiling (SSv4) of fluorescently labeled FACS-isolated cells. Others that produced broad labeling were not analyzed by SSv4. The single-cell transcriptomes were then mapped to the VISp taxonomy as above^9^ (**Figures 7, 8**).

**Figure 7:**
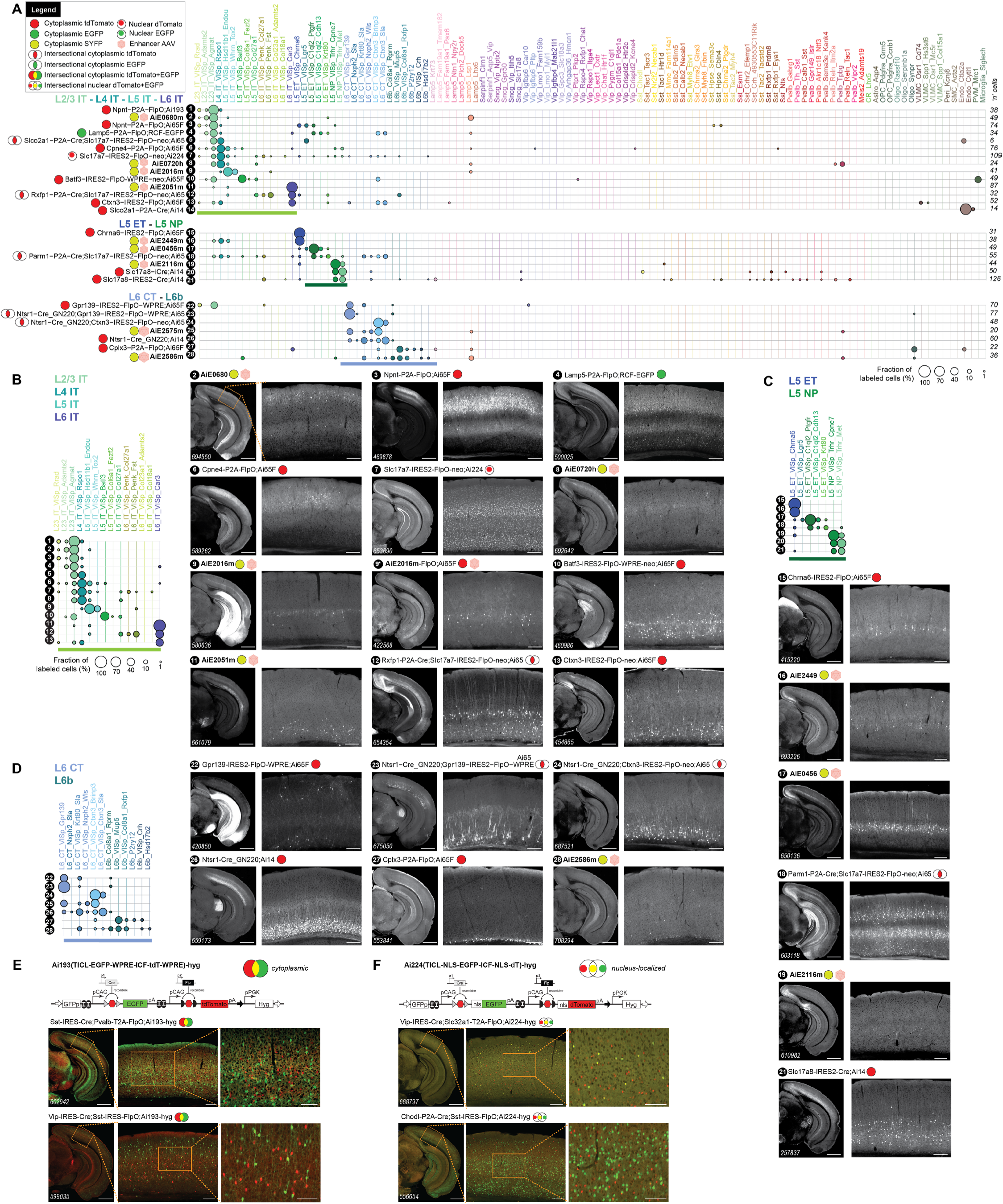
Characterization of new transgenic driver lines, preferentially targeting glutamatergic subclasses and clusters. **A.** scRNA-seq (SSv4) data showing distribution of labeled cells mapped to the mouse VISp taxonomy at the cluster level. 28 different experimental conditions (tools numbered **1-28**) were grouped into panels according to predominant cell types labeled. They may include transgenic drivers and reporters as indicated or may be wild-type animals that received a systemic delivery of enhancer viruses (marked by pink hexagons). **B.** Focused view of tools **1-13** that label IT neurons from Layer 2-3, L4, L5 and L6, including the previously reported enhancer AiE2016m (originally called mscRE16)^22^ expressing SYFP in a wild-type animal (**9**) and driving a FlpO recombinase in Ai65F (**9\***). **C.** Same as in (B) for tools **14-21** that label ET and NP neurons in L5. **D.** Same as in (B) for tools **22-28** that label L6_CT and the L6b neurons. **E.** Schematics and representative sections from STPT data for a new Flp-Cre:AND/OR reporter line, *Ai193* (TICL-EGFP-WPRE-ICF-tdT-WPRE)- hyg. The line was tested in triple transgenic crosses with two recombinase lines. Cells that express the Cre recombinase are EGFP-positive (green) and those that express FlpO are tdTomato-positive (red). Cells that express both appear yellow. **F.** Same as in (E) for a new reporter line, *Ai224* (TICL-NLS-EGFP-ICF-NLS-dT)- hyg, where fluorophores are nucleus-localized. Scale bars: 1.0 and 0.2 mm for full section and expanded view; 0.1 mm for further expanded view in (E) and (F).

Many lines were included into intersectional crosses to examine if additional specificity can be achieved up to the level of single transcriptomic cell-type cluster. The intersectional crosses were performed by first making double recombinase transgenics (Cre and Flp; **Figure 1A**), followed by crossing to a dual “AND” recombinase reporter (e.g., *Ai65* or *RCFL-H2B-GFP*)^69,70^. This crossing scheme prevents unintended recombination and permanent reporter modification in the germline with certain recombinase lines, if they are crossed to a reporter first, and then to the second recombinase^71^. The triple transgenic mice were examined by STPT and SSv4 (**Figures 7, 8**). In most cases, the use of triple transgenics refined reporter expression and resulted in greater specificity (**Table S1**). For comparison, we present equivalent data from enhancer viruses that were RO-injected into either wild-type or reporter mice. In some cases, the viruses were further validated by injecting them into triple transgenic lines that already showed target cell-type labeling (**Figures 8B_29+31_, 8B_30+31_, 8C_39+40_**) and had observed expected overlap between the two labeling patterns.

**Figure 8:**
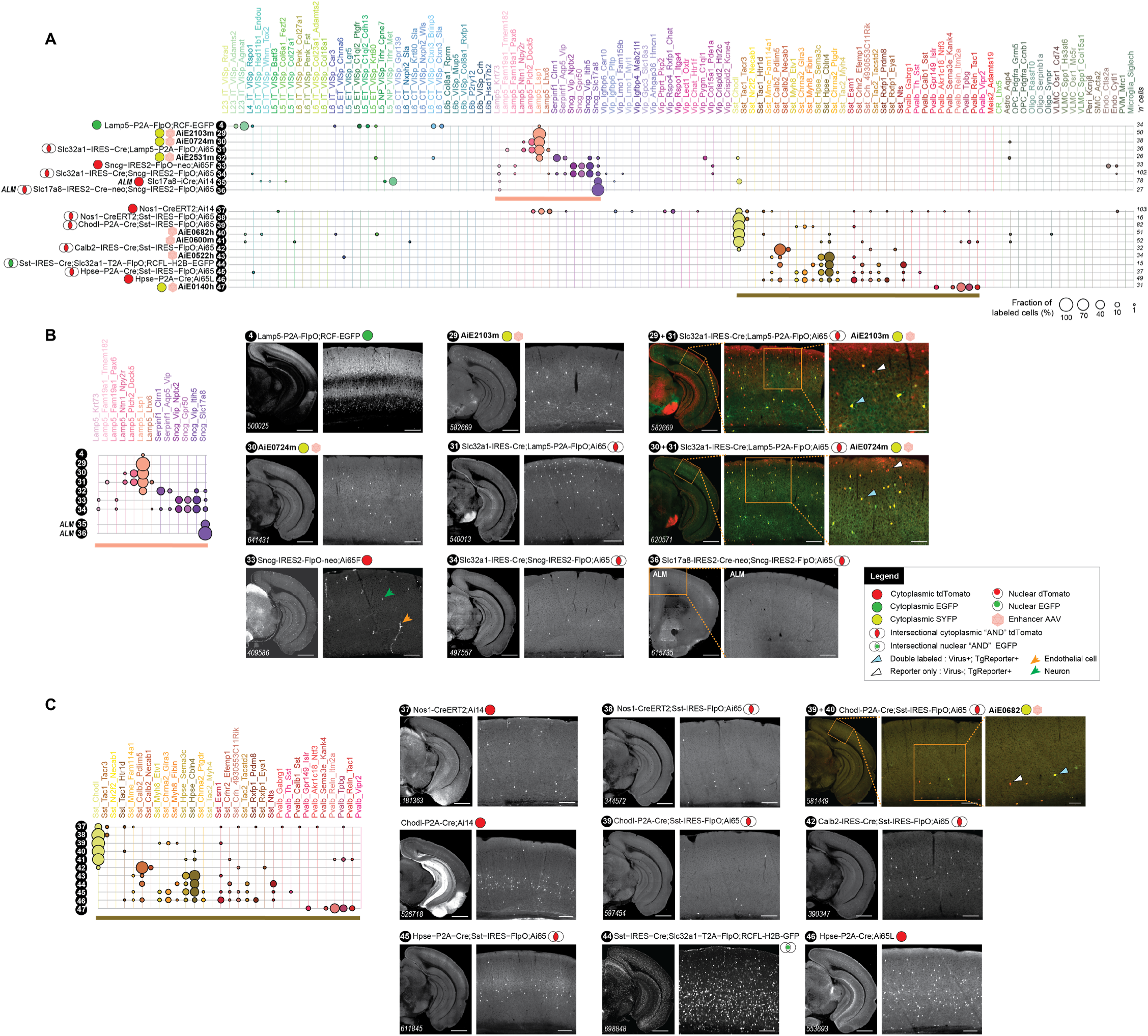
Characterization of new transgenic driver lines, preferentially targeting GABAergic subclasses and clusters. **A.** scRNA-seq (SSv4) data, same as in Figure 7A but for 20 lines (tools numbered **4, 29-34**, **37-47**) targeting GABAergic types grouped into panels according to predominant cell types labeled. **B.** Focused representation of the same data as in (A) for tools labeling clusters within Lamp5 and Sncg GABAergic cortical subclasses. All data (tools numbered **4**, **29-34**) are from VISp, except for tools **35** and **36** that were characterized in cortical area ALM. The tools **29+31** and **30+31** show expression of two different Lamp5-expressing viruses in the triple transgenic mouse that expresses highly specifically only in Lamp5 interneurons (*Slc32a1-IRES-Cre;Lamp5-P2A-FlpO;Ai65* – see tool **31**).**C.** Focused representation of the same data as in (A) for tools **37-47** labeling clusters in Sst and Pvalb GABAergic cortical subclasses. Scale bars: 1.0 and 0.2 mm for full section and expanded view; 0.1 mm for further expanded view for tools **29+31**, **30+31**, **39+40**.

Some cell type taxons, such as the IT glutamatergic subclasses and clusters, are difficult to target with high specificity (**Figure 7A**), perhaps because many markers for IT types tend to be continuously expressed across related IT clusters and subclasses^9,72^. This appears to be true as well for the enhancer viruses that target these populations (**Figure 4B**). A notable exception is the L6_IT_Car3 cluster, which is labeled by several unique markers in addition to being specifically targeted by several enhancer viruses (**Figures 4B**, **7A_11_**, **7B_11_**).

We successfully targeted L5_ET types by viruses at both subclass and cluster level (**Figures 4B**, **S4B**, **7A_16,17_**, **7C_16,17_**). We have also generated a specific transgenic line targeting the *Chrna6* gene that labels the unique L5_ET_Chrna6 cell type (**Figures 7A_15_, 7C_15_**). For the L5_NP subclass, we have previously profiled transgenic lines based on the *Slc17a8* gene (*Slc17a8-iCre* and *Slc17a8-IRES-Cre*)^9,73^. Here, we report a highly specific enhancer virus for the L5_NP subclass (**Figures 4B**, **7A_19_**, **7C_19_**).

For L6_CT and L6b types, we show that a judicious use of crosses can result in nearly single cell-type cluster labeling (**Table S1**). For example, *Ctxn3-IRES2-FlpO* labels an assortment of glutamatergic IT cell types, as well as select L6_CT clusters (**Figures 7A_13_, 7B_13_**). Crossing this line with a L6_CT-specific line, *Ntsr1-Cre_GN220*^74^ (**Figures 7A_26_, 7D_26_**), and then to the intersectional reporter *Ai65*, produces a triple transgenic that precisely labels the only two *Ctxn3*-expressing L6_CT types (L6_CT_VISp_Ctxn3_Sla and L6_CT_VISp_Ctxn3_Brinp3; **Figures 7A_24_, 7B_24_**; **Table S7**). Similarly, we generated a tool that can exhibit a remarkably specific pattern of expression at the level of single L6_CT cluster in a triple transgenic cross. *Gpr139-IRES2-FlpO*, a line based on *Gpr139*, is expressed in several glutamatergic clusters in different layers, including a specific L6_CT cluster. When this line is crossed with *Ntsr1-Cre_GN220* and *Ai65*, we find that it almost exclusively labels only the L6_CT_VISp_Gpr139 cluster (**Figures 7A_22-23_, 7D_22-23_**).

The two new reporters we made, *Ai193*^65^ and *Ai224*^66^ (**Figure S7B**), report Cre “AND/OR” Flp labeling, unlike *Ai65*^70^, which requires both Cre “AND” Flp to express a fluorescent protein (FP, tdTomato in this case). These two lines have separate transcription units that independently report the presence of Cre or Flp, resulting in GFP expression for Cre, tdTomato (or dTomato) expression for Flp, and both FPs when Cre and Flp are present. *Ai193* expresses the FPs in the cytoplasm (**Figure 7E**), whereas *Ai224* expresses them in the nucleus (**Figure 7F**). When compared with previously widely used reporters (*Ai14*, *Ai65* or *Ai65F*; **Figure S8D**), both lines appear to faithfully report Cre and Flp recombinase expression by expressing either one or both marker genes. Note that the localization of GFP in *Ai224*, while predominantly nuclear, is imperfect (**Figure S8F**).

For GABAergic cell types, we selected the *Lamp5*, *Sncg* and *Chodl* genes for targeting. Although *Lamp5* is an excellent marker for the Lamp5 subclass of GABAergic cells, it also labels glutamatergic cells in L2-3, L5 and L6 (**Figures 7A_4_, 7B_4_**, **8A_4_**, **8B_4_**). To exclude expression in glutamatergic cells, we generated a triple transgenic containing the pan-GABAergic line *Slc32a1-IRES-Cre*^75^. We show that *Lamp5-2A-FlpO;Slc32a1-IRES-Cre*;*Ai65* exclusively labels the Lamp5 subclass of GABAergic interneurons in the cortex (**Figures 8A_31_, 8B_31_**). We also find that this subclass can be very specifically labeled by many enhancer viruses (**Figures 8A_29-30_, 8B_29-30_**). Another subclass of MGE-derived GABAergic cells is labeled by the gene *Sncg*. *Sncg-IRES-FlpO;Ai65* labels the *Sncg* population of neurons as well as endothelial cells (**Figures 8A_33_, 8B_33_**). The generation of the triple transgenic *Sncg-IRES-FlpO;Slc32a1-IRES-Cre*;*Ai65* experimental animals excluded the endothelial cells, labeling only the Sncg GABAergic cells (**Figures 8A_34_, 8B_34_**). In addition, we were able to use the Sncg line to label an individual cell type cluster, Sncg_Slc17a8, a unique Sncg type enriched in the frontal cortex^9^. To achieve this, we crossed this line with the *Slc17a8-IRES-Cre* driver line^73^ followed by *Ai65*. We observed exquisitely specific labeling for the Sncg_Slc17a8 cluster, as shown by SSv4 of cells isolated from the Anterior Lateral Motor (ALM) cortex of the triple transgenic *Sncg-IRES-FlpO;Slc17a8-IRES-Cre*;*Ai65* mice (**Figures 8A_35-36_, 8B_36_**). Note that the new Sncg line is useful beyond cortex: when combined with reporter viruses, we used it to characterize the Sncg GABAergic neurons in the hippocampus^76^.

To gain access to the Sst_Chodl cell type cluster, which corresponds to sleep-active *Nos1* and *Tacr1*-expressing cells^77^, we made a Cre line that targets the gene *Chodl* (*Chodl-P2A-Cre*). In initial characterization, we found that this line also labels other cell types (**Figure 8C**), so we generated the triple transgenic *Chodl-P2A-Cre;Sst-IRES-Flpo;Ai65* that exhibits very specific Sst_Chodl cell type labeling (**Figure 8C_39_**). We also discovered two enhancer viruses that quite specifically label this cell type (**Figures 4B,C, 8C_40_**). We note that one of the enhancers (AiE0600m) is located ∼250 kbp from the *Nos1* gene (**Figure 2A**). The previously described *Nos1-CreERT2*^78^ also labels Sst_Chodl, but not cleanly (**Figure 8C_37-38_**). However, *Nos1-CreERT2;Sst-IRES-FlpO;Ai65* quite specifically labels the Sst_Chodl cell type (**Figure 8C_37-38_**).

We tested if similar level of specificity can be achieved for the Sst_Hpse clusters. The *Hpse-P2A-Cre* line labels the Sst_Hpse clusters as well as a few cells of almost every other cluster in the Sst subclass. Although the triple transgenic *Hpse-P2A-Cre*;*Sst-IRES-FlpO*;*Ai65* resulted in an enrichment of the *Hpse*^+^ population, it did not completely eliminate other Sst cell type cluster labeling (**Figure 8C_45-46_**; **Table S1**). Therefore, each cross needs to be tested before assuming that the tools based on the marker genes will provide expected labeling.

We note that the exact components used to create a transgenic line influence the labeling pattern that the line will produce. This has previously been observed with lines targeting the parvalbumin gene (*Pvalb*), where *Pvalb-2A-Cre* expresses in L5_ET types as well as thalamus excitatory cells (thalamocortical projection neurons), whereas the *Pvalb-IRES-Cre* line does not^68^. Consistent with previous observations, we note that compared to IRES, the 2A peptide usually produces broader recombinase and therefore reporter expression. This effect is likely a consequence of 2A fusion better capturing the lower end of the range of marker gene expression than IRES. Similarly, the addition of WPRE increases the overall expression level of the recombinase from the targeted locus. For example, *Gpr139-IRES2-FlpO-WPRE*, is expressed in a broader range of cells including L2-3, compared to both *Gpr139-IRES2-FlpO and Gpr139-IRES2-FlpO-neo* (**Figures S7A_5-6_, S7B_4-6_**).

As observed before^9,79,80^, presence of the Neomycin selection cassette (Neo) can affect transgene expression. Some of the lines we include in this paper were generated by gene targeting without drug selection (the Ngai lines; **Figure S7**), whereas others were generated using antibiotic selection and contained Neo. It is a common practice to remove Neo after the line is established^68^. However, we have observed that in many cases, the expression patterns are different with and without Neo, and these differences may be useful depending on the application^79^. In most cases, the removal of Neo results in an expansion of expression as in the case of *Slc17a8-IRES2-Cre* (**Figure S8B_6-7_**). It does not appear to make a difference for the *Gpr139-IRES2-FlpO* line (**Figure S8B_31-32_**) or for the *Chrna6-IRES2-FlpO line* (**Figure S8B_1-2_**). Note that in the latter case, the addition of WPRE in the *Chrna6-IRES2-FlpO-WPRE-neo* did not appear to change cell type labeling: L5_ET_Chrna6 was still the dominant labeled type, with the same rare L6b cluster (L6b_Col8a1_Rprm) expressing the *Chrna6* mRNA also labeled (**Figure S8B_1-3_**)^8,9^.

In this study, we used SSv4 to evaluate the expression pattern of transgenic lines. However, it is important to compare, whenever possible, the SSv4 data with other modalities to confirm labeling patterns. For example, the SSv4 data for *Npnt-2A-FlpO;Ai65* show cells from both L2-3 and L5 whereas that for *Npnt-2A-FlpO;Ai193* only includes L2-3 cells implying *Ai193* only labels a subset of cells expressing Npnt. However, consistent with *in situ* hybridization results (https://mouse.brain-map.org/experiment/show/71670677; **Figure S8C**)^81^, *Npnt-2A-FlpO;Ai65F* and *Npnt-2A-FlpO;Ai193* show identical expression patterns by STPT with labeling of both L2-3 and L5 cells (**Figure S8C_7-8_**). This discrepancy can be explained by our previous report that some cell types (specifically cortical Pvalb and L5 ET cells) are sensitive to FACS isolation, and survive the process with variable success experiment-to-experiment^8,9^. Therefore, the absence of L5_ET cells in some SSv4 experiments (**Figure S8A_9-10_**) is likely an effect of FACS, and not differential reporter sensitivity.

We sought to further explore the effect of reporter line choice by comparing labeling patterns in crosses where the same driver lines were crossed with previously characterized reporters (*Ai14* and *Ai65/Ai65F*) and the new AND/OR reporters (*Ai193* and *Ai224*). In every case examined, we found that there was no discernable difference between the patterns of labelled cells. For example, triple-transgenic animals obtained by crossing *Chodl-P2A-Cre;Sst-IRES-FlpO* with *Ai65* or *Ai224* show a similar number and distribution of labelled cells (red cells in the *Ai65* cross and yellow-double labelled cells in the *Ai224* cross; **Figure S8D**). Likewise, the labelling patterns in double transgenic crosses of *Sst-IRES-FlpO* or *Chodl-P2A-Cre* with *Ai224, Ai14*, *RCL-H2B-EGFP*^69^ or *Ai193,* show that the labeling pattern, at least for these reporters, depends on the recombinase lines employed (**Figure S8D**).

Finally, we expected that the knock-in recombinase-based labeling may be more inclusive of all cells that express a certain marker gene compared to enhancer AAVs. However, we show at least one instance where the enhancer virus faithfully captures the expression pattern of a marker gene while the knock-in recombinase line-based labeling is incomplete. *Cplx3* is a marker gene for L6b neurons as shown by scRNA-seq^9,13^ and RNA *in situ* hybridization^81^ (https://mouse.brain-map.org/experiment/show/70928340; **Figure S8E**). The expression pattern for AiE2359m, as well other L6b enhancers we discovered, mirrors *Cplx3* expression whereas the transgenic cross, *Cplx3-P2A-FlpO;Ai193* excludes *Cplx3+* cells in the lateral entorhinal area, and L1 interneurons, which also express this gene^82^. This effect is independent of the reporter line as we also observe it with a different reporter in *Cplx3-P2A-FlpO;Ai65* (**Figure S8E**).

In summary, we provide a wealth of characterization data for the tools we created and underline the importance of careful evaluation of each tool or a combination of tools for the intended purpose. We strongly suggest evaluation of each tool/tool combination with more than one data modality, as some modalities have modality-specific pitfalls that can be highlighted with a different modality.

## Discussion

The definition of cell types and their taxonomy in the central nervous system and genetic access to individual types are essential for our understanding of how they contribute to nervous system function and dysfunction^1,3,11,83^. We utilized single-cell transcriptomic and epigenomic data to identify marker genes for the generation of new transgenic lines, as well as putative enhancers for the creation of enhancer AAVs.

Currently, the use of transgenic mouse lines is the main approach for gaining access to molecularly identified cellular populations^8,67,84,85^. Transgenic mouse lines are created by inserting exogenous DNA (e.g., fluorophore, recombinase or transcription factor) into the mouse genome. A subset of transgenic mouse lines, the knock-in lines, are created by inserting a single copy of exogenous DNA, at a specific position, frequently within a marker gene in the mouse genome^67,86^. All the transgenic driver lines described in this paper were generated by knocking-in a recombinase into the endogenous gene locus. This contrasts with randomly integrated transgenes where the endogenous elements have been taken out of context and depending on the size/regulatory elements included, copy number and insertion site, dramatic variations in labeling patterns may occur^74,87^ or may even behave as enhancer traps^88^.

The knock-in approach takes advantage of the endogenous genomic regulatory elements to enable selective expression of various transgenes for cell labeling, monitoring and/or perturbation, and frequently produces expected cell-type labeling. However the efficiency, specificity, and strength of cell type labeling by transgenes generated in the same locus can vary depending on the exact components inserted^68,89^, and we report several examples of this phenomenon in this study. In a number of cases, the driver lines labeled unintended cell types in which the marker gene expression was weak or not observed in adult mice, which is important to note especially if functional reporters are to be driven by them^90^. In order to restrict recombination to the target population, we generated triple transgenic mice using two separate driver lines, whose recombination patterns intersect only in the population of choice, with a dual reporter line requiring the presence of both recombinases for its expression^24,69,70,91^.

Generation, validation, and maintenance of transgenic lines is expensive and laborious. Transgenes are integrated into the genome and although methods for their modification once established have become recently available^92^, these are not widely utilized. Moreover, site-specific transgenesis in other mammalian species is difficult and costly^93,94^ Therefore, it is not practical to rely on transgenic lines for all experiments. The use of enhancer AAVs^17,18,22,30,95^ and other viral^32,96^ or non-viral technologies^97^ that do not require germline modification can potentially circumvent all these obstacles. However, there are caveats to be considered when applying enhancer AAVs: 1) even for the same vector, the degree of specificity and expression strength heavily depend, among other factors, on the concentration, delivery method, and viral serotype, which can lead to larger variability within and across experiments than with mouse transgenes; 2) access to some populations can be more difficult, due to their decreased susceptibility to viral transduction (e.g., microglia)^64^. For any untested crosses or combinations of transgene and virus, we advise the user to characterize them before assuming specificity based on the marker genes or enhancers used.

In this study, we provide a detailed and comprehensive report on the development, screening and use of enhancer AAVs in the context of the mouse cortex. We demonstrate that, at the subclass level of cell-type resolution, the mammalian species of origin appears to have little effect on enhancer performance, suggesting that many of the enhancer sequences described here could be useful in other mammalian species, even if their orthologs are not present in the species of interest. To evaluate enhancer specificity at a transcriptomic cell type (cluster) resolution, we performed scRNA-seq on isolated cells labeled by individual enhancer AAVs. We confirmed that many enhancer AAVs designated as ‘promising’ in primary screen showed high specificity for a single cell type taxon either at the subclass or cluster level (**Figure 4, Figure S4A**). We also found that the SYFP mRNA expression was significantly lower when driven by enhancer AAVs compared to the pan-neuronal hSyn1 promoter/enhancer, which could make enhancers insufficiently strong to drive functional cargo such as effectors and indicators. However, we show that in many cases, this limitation can be overcome by optimization of enhancers through core bashing and concatenation (**Figure 5**), expression of a recombinase (**Figure 6**) or by delivering viral vectors using alternative routes, such as ICV or stereotaxically (**Figures S5, S6**). We demonstrate that all these approaches can lead to a marked increase in cargo expression, often with tolerable specificity loss.

The tools reported here, as well as the scaled and standardized process used to create and evaluate them, provide an unprecedented resource that should enable diverse experimental strategies for understanding mammalian cortex function, including access to many previously inaccessible cortical types. We are in the process of making all materials, data and metadata associated with the study publicly available. Moreover, in an associated study^98^, our standardized experimental process for enhancer evaluation described here has advanced our understanding of the basic features underlining enhancer performance.

## Materials and Methods

### Animals

Mice were housed in the Allen Institute Vivarium and all animal procedures were conducted in accordance with the Institutional Animal Care and Use Committee (IACUC) protocols 1508, 1802, 1806, 2105, and 2406, with no more than five animals per cage, maintained on a 14/10 h day/night cycle, with food and water provided *ad libitum*. Both male and female mice were used for experiments and the minimal number of animals were used for each experimental group. Animals with anophthalmia or microphthalmia were excluded from experiments. Animals were maintained on a C57BL/6J genetic background. At the University of California, Berkeley, experiments with mice were conducted under the campus’s Animal Care and Use Committee Animal Use Protocol # AUP-2016-08-9032. See **Table S2** for a full list of all transgenic mouse lines included in this study.

### 10x Genomics snMultiome data generation

Mice were anaesthetized with 2.5–3% isoflurane and transcardially perfused with cold, pH 7.4 HEPES buffer containing 110 mM NaCl, 10 mM HEPES, 25 mM glucose, 75 mM sucrose, 7.5 mM MgCl_2_, and 2.5 mM KCl. Mice were anesthetized with 2.5-3.5% isoflurane. Following perfusion, brains were isolated quickly, and frozen for 2 minutes in liquid nitrogen vapor. Frozen brain samples were sectioned on a cryostat to obtain 300 µm sections. Micro knives were used to microdissect the area of interest according to the Allen reference atlas. Images were collected pre- and post-microdissection to document which brain regions were profiled. Dissectates were placed in 12-well plates in the cryostat after collection.

Following dissection, nuclei were isolated with the RAISINs protocol (RNA-seq for Profiling Intact Nuclei with Ribosome-bound mRNA) # PF0334^99^ developed based on a previously published protocol^100^. Briefly, tissue samples were placed in CST buffer and single cell suspensions were obtained by chopping tissue using spring scissors for 10 minutes in the buffer. Cell suspensions were centrifuged 500 rcf and resuspended in lysis buffer. Nuclei were counted, resuspended, and processed according to the 10x multiome protocol from 10x Genomics. Short-read sequencing was done with Illumina. Fastq generation and alignment to mm10 was done with Cell Ranger ARC (version 2.0.0). Downstream data analysis was performed with scanpy (version 1.9.8).

Human MTG data were collected by subsetting healthy control data from the SEA-AD study^28^. Briefly, brain specimens were donated for research to the University of Washington BioRepository and Integrated Neuropathology (BRaIN) laboratory from participants in the Adult Changes in Thought (ACT) study and the University of Washington Alzheimer’s Disease Research Center (ADRC). 10x multiome library preparation was performed as per the 10x multiome protocol (10x Genomics). Sequencing was performed using a NovaSeq 6000, using either a NovaSeq-X or S4 flow cell. Fastq generation and data alignment to GRCh38 was done with Cell Ranger ARC. Downstream data processing was done using the scanpy python package (version 1.9.1).

### snMultiome: Mouse snRNA-seq data analysis

Cell clustering and filtering was performed using the standard scanpy workflow. Following this, individual datasets across cortical regions were integrated using SCVI, using individual donors as the batch key. Final cell type annotations for both species were derived from de novo clusters (scanpy). Low quality clusters (containing doublets) were removed after clustering. This was done iteratively. Low quality cells (fewer than 2,000 genes detected for neurons, and fewer than 1,000 genes detected for non-neuronal clusters) were removed. For each de novo cluster, the predominant cell type was used to label the cluster with the appropriate cross-species subclass label. Known marker genes as identified from the Tasic et al 2018 cortical taxonomy were used to check cell type identities. Gene expression data were plotted using scanpy.pl.umap.

### snMultiome: snATAC-seq data analysis

snMultiomic ATACseq data were analyzed using ArchR^40^. Pseudobulk coverage data were used for identification of peaks in VISp subclasses using Macs2 as implemented by ArchR (version 1.0.2) ^40,41^. Statistically significant peaks were identified using ArchR. Bigwig files were used for enhancer selection, data visualization and for determination of accessibility at putative enhancer sites. For downstream analyses, enhancer accessibility was obtained by summing ATAC signal in subclass-specific bigwig files in all bins overlapping enhancer regions. This was done using the GenomicRanges package in R. Accessibility was scaled across subclasses to obtain z-scores for enhancer accessibility at each enhancer site. This measure was used to characterize enhancer target populations in an unbiased way.

### Cross-species analysis

Enhancer sequences were obtained from mm10 using genomic coordinates using the Biostrings package in R. Mouse sequences were lifted over to hg38 and human sequences were lifted over to mm10. BLAST was performed in a similar manner, using rBLAST using the following arguments: word size = 10; reward = 2; penalty = 3; gapopen = 5; gapextend = 2; dust = no; soft masking = false. Enhancers that returned a match using both liftover and BLAST were considered to have conserved sequence overall. Alignment statistics from BLAST output were retained for each enhancer and were used to calculate a percent of bases in the original enhancer sequences that overlapped with the aligned region. Accessibility correlation between mouse and human was done for each enhancer, using signal from bigwig files across all cortical subclasses.

### Enhancer cloning

Short enhancer sequences were nominated from the mouse or human genome based on selective chromatin accessibility in either of the cortical subclasses, with preference for sequences found in proximity to marker genes for that subclass. These sequences were subsequently amplified from purified genomic material of wt C57BL6/J mice, cloned into an AAV2-minPromoter-SYFP2-WPRE-bGH backbone upstream of the minimal promoter (BetaGlobin, CMV or Rho) along with a bovine growth hormone polyA (BGHpA), and a woodchuck post-transcriptional regulatory element (WPRE or WPRE3) ^101^. Cloning was carried out either using the Gibson assembly method (NEB; Catalog# E2621L) or restriction digestion and ligation. The recombinant plasmids were verified with Sanger sequencing. Select plasmids have been submitted or are in the process of being submitted to Addgene for distribution. See **Table S3** for a full list of all plasmids included in this study.

### AAV Production

Verified plasmids were packaged into AAV vectors by transient transfection of HEK 293T cells (ATCC CRL-11268). The cells were seeded at 2 × 10^7^ cells per 15-cm dish to achieve ∼70% to 80% confluency before transfection. Cells were maintained in DMEM (Thermo Fisher Scientific Cat#1995-065) with 10% Fetal Bovine Serum (FBS; VWR Cat#89510-184) and 1% Antibiotic-Antimycotic solution (Sigma Cat#A5955). Each enhancer AAV vector was mixed with pAdenoHelper and PHP.eB rep-cap plasmids in a ratio of 30:15:15 µg in 1.35 ml of OptiMem (Thermo Fisher Scientific Cat#51985-034), which was then supplemented with 150 µl of 1 mg/mL Polyethylenimine (PEI; Polysciences Cat#23966), incubated for 10 minutes and then added to a single 15 cm plate of fully confluent cells. Twenty-four hours post transfection, the cell medium was changed to media containing 1% FBS and 1% Antibiotic-Antimycotic in DMEM and 72 hours later, cells were harvested into a 50 ml-tube, subjected to three 20-minute long freeze-thaw cycles to lyse cells and release adeno-associated virus (AAV) particles, and the resulting lysate was incubated with benzonase for an additional 30 minutes (Sigma-Aldrich Cat#E1014) at 37°C, to remove non-encapsidated nucleic acids. The crude AAV-containing suspension was centrifuged at 3000*xg* for 10 minutes to remove residual cell debris, and the supernatant was concentrated using an Amicon Ultra-15 centrifugal filter (Sigma Cat # UFC910024) by centrifuging at 3000*xg*, until the volume was reduced to below 150 µl. This crude AAV prep was then aliquoted and kept at −80°C until use. ^102–104^.

For vectors intended for stereotaxic delivery, the transfected cells were pelleted and resuspended in 1 ml lysis buffer containing 150 mM NaCl and 50 mM TrisHCl (pH=8), instead of in their growth media, and following the freeze-thaw cycles and Benzonase treatment, the lysate was passed through a 0.22 µm filter, to remove any large debris which might clog the capillary.

### AAV titer determination with dd-PCR

For measuring virus titers, we used ddPCR (Bio Rad; QX 200 Droplet Digital PCR System). We used primers against AAV2 ITR for amplification. Seven serial dilutions with the factor of 10 ranging from 2.5×10^-2^ to 2.5×10^-8^ were used for the measurement. Serial dilutions of 2.5×10^-5^ to 2.5×10^-8^ were used for fitting the linear dynamic range. Viral titer was calculated by averaging virus concentration of two dilutions within the linear dynamic range. A positive control of a known viral titer, and a negative control with no virus was also run along with all the samples.

### Retroorbital (RO), Intracerebroventricular (ICV) and Stereotaxic (STX) virus injections

RO injections were performed according to a previously described protocol^44^: Male and female C57BL/6J mice, aged P27–P33, were anesthetized using isoflurane. Each mouse received an injection of 5×10^11^ genome copies (GC) diluted to 90 µl with PBS, administered into the retro-orbital sinus.

Intracerebroventricular (ICV) injections were performed according to the previously described protocol^105^: P0-2 mouse pups were anaesthetized by placing them on aluminum foil on ice and injected with 5 x 10^10^ GC diluted to 5 µl with PBS, targeting the lateral ventricles. The injection was made 1 mm lateral to the midline and approximately 1 mm posterior to bregma. Mice were euthanized after a four-week incubation period.

For stereotaxic (STX) injections, male and female C57BL/6J mice, aged P45–P90, were anesthetized with isoflurane before injecting with filtered PHP.eB-pseudotyped AAV unilaterally or bilaterally into the primary visual cortex (VISp) using the following coordinates (in mm): anterior/posterior (A/P) −3, medial/lateral (M/L) ±2, and dorsal/ventral (D/V) 0.45/0.65. A total volume of 300 nl containing 1.5×10^9^ GC/ml virus was delivered at a rate of 50 nl per pulse with Nanoinject II. Before incision, the animal was injected with Bupivacaine (2-6 mg/kg) and post injection, the animal was injected with ketofen (2-5 mg/kg) and Lactated Ringer’s Solution; LRS (up to 1 ml) to provide analgesia. Mice that underwent STX injections were euthanized after 25-31 days, transcardially perfused, and the brains were dissected for further analysis.

### Brain tissue preparation and image acquisition

Mice were anaesthetized with isoflurane and perfused transcardially with 10 ml of 0.9% saline, followed by 50 ml of 4% PFA. The brain was removed, bisected sagittally along the midline, placed in 4% PFA overnight and subsequently moved to a 30% sucrose in PBS solution until sectioning. From the left hemisphere, 30 µm sections were obtained along the entire mediolateral axis using a microtome. Five sections, roughly 0.5, 1, 1.5, 2.3 and 3.5 mm from the midline, were collected, stained by DAPI and/or PI to label nuclei and cellular RNA, and after drying for 24 hours at 37 °C, mounted on a barcoded slide. Once the mounting medium hardened, the slides were scanned with Aperio VERSA Brightfield epifluorescence microscope (Leica) in the UV, green, and red channels, illuminated with a metal halide lamp. After passing QC, digitized images were analyzed by manual scoring.

### Primary screen scoring

Each enhancer vector was scored based on the labeling pattern it produced in the neocortex. First, each region of the brain where labeling of cell somata was observed, was manually scored based on the labeling brightness and density, classifying each into either low or high. In addition, we created 11 categories of different cell populations within the neocortex, which could be visually distinguished one from the other, and whenever cortical labeling was observed, in one or more of these populations, each was individually evaluated based on its own brightness and density. Whereas brightness was classified based on whether the labeling was stronger or weaker than the common brightness observed across all experiments, density was evaluated based on the expected density of cells for each of the scored regions or populations, using the nuclear markers as reference. To determine target specificity, we aligned each target cell population with the labeled population which best matches it’s known anatomical location, distribution, and morphological characteristics. We determined an enhancer to be “On-target” if the target population aligned with the labeled population, “Mixed” if labeling was observed in populations other populations, in addition to the target one, “Off-target” if labeling was observed exclusively in population/s other than the target one, and “no labeling” if no labeling was observed in the neocortex, regardless of whether labeling was observed in other brain regions.

### SMART-Seq v4 sample preparation and analysis

Sample preparation for SMART-Seq was performed using the SMART-Seq v4 kit (Takara Cat#634894) as described previously^9^. In brief, single cells were sorted into 8-well strips containing SMART-Seq lysis buffer with RNase inhibitor (0.17 U/µL; Takara Cat#ST0764) and were immediately frozen on dry ice for storage at −80°C. SMART-Seq reagents were used for reverse transcription and cDNA amplification. Samples were tagmented and indexed using a NexteraXT DNA Library Preparation kit (Illumina Cat#FC-131-1096) with NexteraXT Index Kit V2 Set A (Illumina Cat#FC-131-2001) according to manufacturer’s instructions except for decreases in volumes of all reagents, including cDNA, to 0.4 x recommended volume. Full documentation for the scRNA-seq procedure is available in the ‘Documentation’ section of the Allen Institute data portal at http://celltypes.brain-map.org/. Samples were sequenced on an Illumina HiSeq 2500 as 50 bp paired end reads. Reads were aligned to GRCm38 (mm10) using STAR v2.5.3^106^ with the parameter “twopassMode,” and exonic read counts were quantified using the GenomicRanges package for R as described in Tasic et al. (2018). To determine the corresponding cell type for each scRNA-seq dataset, we utilized the scrattch.hicat package for R^72^ (https://github.com/AllenInstitute/scrattch.hicat). We selected marker genes that distinguished each cluster, then used this panel of genes in a bootstrapped centroid classifier which performed 100 rounds of correlation using 80% of the marker panel selected at random in each round. For plotting, we retained only cells that were assigned to the same cluster in ≥ 80 of 100 rounds. Cells that did not map to the taxonomy confidently were excluded from analysis and further data processing. Mapping results and scRNA-seq sample metadata, including the most-frequently assigned cell type and the fraction of times each cell was assigned to that type, are included in supplemental data.

For experiments involving enhancer AAVs, mice were injected retro-orbitally with the indicated AAV. One month post injection, individual cells were FACS-isolated from cortical regions. In most cases this was mouse visual cortex, cells were also collected from claustrum. In most cases, all VISp layers were isolated prior to FACS. In others, two or more collections were made using an “upper” (layers 1-4) and a “lower” (layers 5-6) dissection strategy, and the data were pooled. In summary, for transgenic lines, cells were summed across all layer collections, which may include single layers or combinations. This may contribute to biases in cell-type compositions reported.

### Generation of driver lines at UC, Berkeley

To generate mouse lines bearing in-frame genomic insertions of P2A-FlpO or P2A-Cre, we engineered double-strand breaks at the stop codons of the targeted genes using ribonucleoprotein (RNP) complexes composed of SpCas9-NLS protein and in vitro transcribed sgRNA for the following gene targetings:

Cplx3-P2A-FlpO (sgRNA: GGGGAAGTGGTCACATGATA);

Cpne-P2A-FlpO (sgRNA: ATATGAATCGTCCCGGACAC);

Npnt-P2A-FlpO (sgRNA: GATGATGTGAGCTTGAAAAG);

Lamp5-P2A-FlpO (sgRNA: CCAGTACAAGCACATGGGCT);

Chodl-P2A-Cre (sgRNA: ATGGAGGTATAATAATGAAC);

Hpse-P2A-Cre (sgRNA: TCATATACAAGCAGCGATTT);

Parm1-P2A-Cre (sgRNA: CGTTAAGAGTCATCGTAGAG);

Rxfp1-P2A-Cre (sgRNA: ACTCAATTCTTATTCGTAAC);

Slco2a1-P2A-Cre (sgRNA: CAGTCTGCAGGAGAATGCCT).

The RNP complexes were nucleofected into 10^6^ v6.5 mouse embryonic stem cells (C57/BL6;129/sv; a gift from R. Jaenisch) along with repair constructs in which P2A-FlpO or P2A-Cre was flanked with homologous sequences 5’ and 3’ to the target site, thereby enabling homology-directed repair. Colonies grown from transfected cells were screened by PCR for successful integration; proper insertion of the transgene was confirmed by DNA sequencing. Cell lines with normal karyotypes were aggregated with albino morulae and implanted into pseudopregnant females to produce germ line competent chimeric founders, which were then bred to C57BL/6J mice for further analysis.

### Generation of driver lines at Allen Institute

Knock-in driver lines contained components that were previously described^67,68,70^. Targeting of the transgene cassettes into an endogenous gene locus was accomplished via CRISPR/Cas9-mediated genome editing using circularized targeting vector in combination with a gene-specific guide vector (Addgene plasmid #42230). The129S6B6F1 ES cell line, G4, was utilized directly for driver line targeting. Targeted ES cell clones were subject to standard antibiotic selection and correctly targeted ES cells were identified using standard screening approaches (PCR, qPCR, and Southern blots) and injected into blastocysts to obtain chimeras and subsequent germline transmission. The resulting mice were crossed to the Rosa26-PhiC31 mice (JAX Stock # 007743) to delete the pPGK-neo selection marker cassette, and then backcrossed to C57BL/6J mice and maintained in C57BL/6J congenic background. Only mice heterozygous for both reporter and driver transgenes were used for experiments.

### Generation of TIGRE3.0 reporter lines

To target multiple transgene expression units into the *TIGRE* locus we employed a recombinase-mediated cassette exchange (RMCE) strategy similar to that previously described^70^, but instead of using Flp recombinase for targeting, we used Bxb1 integrase^107^ so that Flp recombinase could later be used for transgene expression control. A new landing pad mouse embryonic stem (ES) cell line was generated by taking the 129S6B6F1 cell line, G4^108^, and engineering it to contain the components from 5’ to 3’ Bxb1 AttP-PhiC31 AttB-PGK promoter-gb2 promoter-Neomycin gene-PGK polyA-Bxb1 AttP-splice acceptor-3’ partial hygromycin gene-SV40 polyA-PhiC31 AttP within the *TIGRE* genomic region. Southern blot, qPCR and junctional PCR analyses were performed on genomic DNA (gDNA) samples from modified ES cell clones to confirm proper targeting, copy number, and orientation of the components within the *TIGRE* locus. A Bxb1-compatible targeting vector with three independent and conditional expression units was then generated by standard molecular cloning techniques. The vector contained the following components from 5’ to 3’: gb2 promoter-Neo gene-Bxb1 AttB-partial GFP-2X HS4 Insulators-CAG promoter-LoxP-stop-LoxP-EGFP-WPRE-BGH polyA-2X HS4 Insulators-CAG promoter-FRT-stop-FRT-mOrange2-HA-WPRE-BGH polyA-PhiC31 AttB-WPRE-BGH polyA-2X HS4 Insulators-CAG-nox-stop-nox-mKate2-P2A-WPRE-PGK polyA-PhiC31 AttB-PGK promoter-5’ hygromycin gene-splice donor-Bxb1 AttB. The sequence and integrity of the targeting vector was confirmed by Sanger sequencing, restriction digests and *in vitro* testing performed in HEK293T cells. The targeting vector (30 µg of DNA) was then co-electroporated with a plasmid containing a mouse codon optimized Bxb1 gene under the control of the cytomegalovirus (CMV) promoter (100 µg of DNA) into the Bxb1-landing pad ES cell line and following hygromycin drug selection at 100-150 µg/ml for 5 days, monoclonal populations of cells were hand-picked and expanded. Genomic DNA was prepared from the modified ES cell clones using a kit (Zymo Research Cat#D4071) and it was screened by qPCR and junctional PCR assays to confirm proper targeting into the *TIGRE* locus. Correctly targeted clones were injected into fertilized blastocysts at the University of Washington Transgenic Research Program (TRP) core to generate high percentage chimeras and then the chimeras were imported to the Institute, bred to C57BL/6J mice to produce F1 heterozygous reporter mice, and subsequently maintained in a C57BL/6J congenic background.

### Statistical analyses

All values were shown as mean and error bars as ± SEM in Figures and reported as mean ± SD in the main text. Statistical significance was tested with a 1-way ANOVA, followed a Tukey test for post-hoc comparisons, or by the Chi square test for analysis of differences in group proportions. All p-values reported were corrected for multiple comparisons using the Bonferroni correction. All calculations were performed in Microsoft Excel or R. Statistical differences with p < 0.05 were considered significant. In Figures, a single asterisk (∗), double asterisks (**), and triple asterisks (***) indicate p < 0.05, p < 0.01, and p < 0.001, respectively.

## Supporting information

Table_S1_New_transgenic_crosses

Table_S2_Transgenic_lines

Table_S3_Plasmids_and_enhancers

Table_S4_Enhancers

Table_S5_HOF_reagents

## Data Availability

Mouse 10x Multiome data and single-cell RNA-seq data are available at the Neuroscience Multi-omic Archive (NeMO):

https://data.nemoarchive.org/biccn/grant/u19_zeng/zeng/multimodal/sncell/10xMultiome_RNAseq/mouse/raw https://data.nemoarchive.org/biccn/grant/u19_zeng/zeng/multimodal/sncell/10xMultiome_ATACseq/mouse/raw

SSv4 scRNA-seq data are available at NeMO: https://data.nemoarchive.org/biccn/grant/rf1_tasic/tasic/transcriptome/scell/SSv4/mouse/raw/

Primary screen epifluorescence and serial-two-photon tomography (STPT) data are available at the Brain Image Library (BIL): https://doi.org/10.35077/g.1162

Primary screen epifluorescence data are available at BIL: https://download.brainimagelibrary.org/4e/fa/4efaa61008dfb900/

Serial-two-photon tomography (STPT) data are available at BIL: https://download.brainimagelibrary.org/bc/59/bc59278fe0669df7/

## Code Availability

Analysis methods used in this manuscript include SCVI (https://github.com/scverse/scvi-tools) and Scanpy (https://github.com/scverse/scanpy) for data processing integration and visualization, ArchR for ATAC-seq analyses (https://github.com/GreenleafLab/ArchR), as well as scrattch.hicat and scrattch.bigcat for cell type mapping and identification (https://github.com/AllenInstitute/scrattch.hicat). Plots were generated using R packages ggplot2, dendextend, which are available from both CRAN and github. Additional code used for data processing and analysis in this manuscript will be made available upon request.

## Author contributions

***Conceptualization***, T.L.D., H.Z., B.P.L., J.N., J.T.T., B.Tasic; ***Methodology - Transgenic lines***, S.N., T.L.D., D.A.S., S.A., T.B., J.Bendrick, J. Bowlus, G.B., B.C., R.K.C., M.D., C.Halterman, O.H., C.Huang, R.L., A.Y.L., M.L., G.H.L., E.L., D.Rette, J.P.S., L.Shulga, L.Siverts, E.S., M.W., T.W., B.W., M.R., K.R., H.Z., B.Tasic; ***Methodology - Viral vectors***, M.H., T.L.D., J.K.M., R.A.M., X.O., J.R.R., N.Donadio, L.G., N.G., F.H., S.H., S.K., E.K., R.K., G.H.L., D.N., K.S., C.T., N.W., T.Z., S.Y., B.P.L., J.T.T., B.Tasic; ***Investigation***, Y.B., M.H., S.N., T.L.D., D.D., J.K.M., M.J.T., J.R.R., A.A., S.B., J.Bendrick, D.B., K.B., T.C., A.B.C., M.C., N.Dotson, T.E., A.G., J.Gloe, L.G., F.H., W.H., S.H., Z.J., M.K., G.H.L., N.L., J.M., R.M., E.M., K.N., V.O., A.Oyama, T.P., E.P., C.A.P., L.P., S.R., C.R., A.R., N.S., A.R.S., M.T., C.T., A.T., N.W., J.A., N.Dee, K.A.S., H.Z., B.P.L., J.T.T., B.Tasic; ***Resources***, S.M.; ***Formal Analysis***, Y.B., M.H., S.N., T.L.D., D.D., A.Oster, R.C., S.C., M.Gabitto, J.G., J.Goldy, B.B.G., L.G., A.H., N.J., B.K., C.L., N.L., E.M., D.Rocha, A.S., S.Somasundaram, C.T.J.V., M.E.W., Z.Y., B.P.L., J.T.T., B.Tasic; ***Data curation***, Y.B., M.H., S.N., D.D., S.W.W., A.Oster, J.R.R., N.G., R.E.A.S., B.P.L., J.T.T.; ***Project Administration***, A.Oster, T.M., R.E.A.S., S.M.S., B.Thyagarajan; ***Visualization***, Y.B., M.H., S.N., D.D., B.Tasic; ***Supervision***, Y.B., S.N., T.L.D., M.J.T., T.E.B., T.C., A.B.C., M.Gasperini, D.H., R.L., T.M., L.N., L.P., C.T.J.V., Z.Y., N.Dee, M.R., K.R., S.M., S.M.S., K.A.S., L.E., J.W., S.Y., E.S.L., H.Z., B.P.L., J.N., J.T.T., B.Tasic; ***Writing – Original Draft***, Y.B., M.H., S.N., D.D., B.Tasic; ***Writing – Review & Editing***, Y.B., M.H., S.N., D.D., S.W.W., B.P.L., J.T.T., B.Tasic; ***Funding Acquisition***, E.S.L., H.Z., B.P.L., J.N., J.T.T., B.Tasic.

## Acknowledgments

We are grateful to the In Vivo Sciences, Molecular Biology, Histology, Imaging, and Neurosurgery & Behavior teams at the Allen Institute for their technical support. We thank Robert Hunter for coordinating transgenic mouse production. We thank Bing Ren and colleagues for open access sharing of the CATlas resource for mouse brain snATAC-seq data. This work was funded by the Allen Institute for Brain Science in addition to the NIH grants U19MH114830 to H.Z., RF1MH121274 to B.T., and RF1MH114126 and UG3MH120095 to E.S.L., J.T., and B.P.L. The authors wish to thank the Allen Institute founder, Paul G. Allen, for his vision, encouragement, and support.

## Declaration of generative AI and AI-assisted technologies in the writing process

During the preparation of this work the authors used Microsoft Copilot in order to improve readability. After using this tool/service, the authors reviewed and edited the content as needed and take full responsibility for the content of the published article.

---

**Figure S1:**
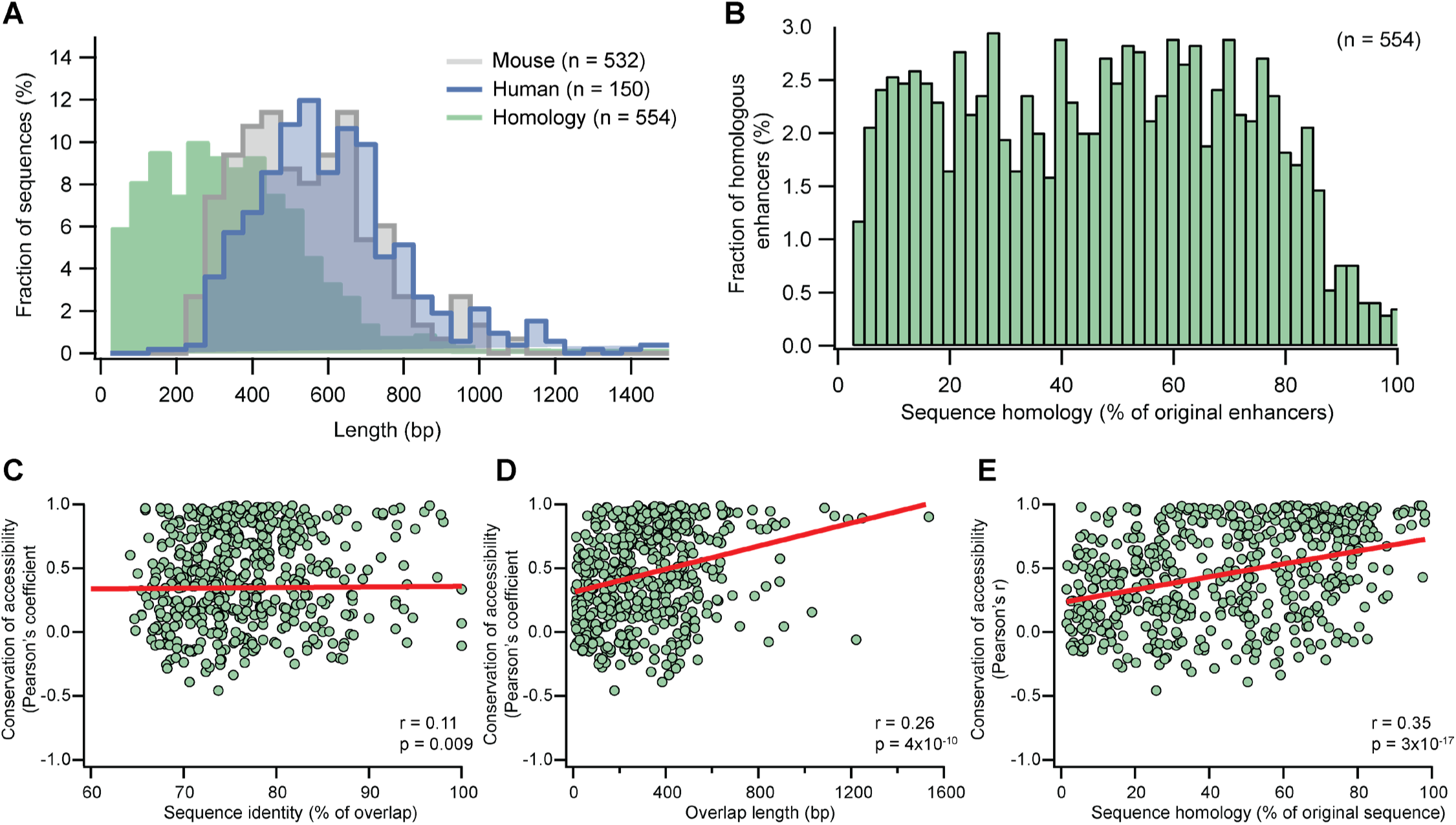
Analysis of sequence alignment across species. **A.** Histograms showing the proportion of mouse (grey) and human (blue) sequences, according to their length along with a histogram showing the length distribution of the liftover sequences (full green). Bin size = 50 bp. **B.** Histogram of the sequence homology, calculated as the total number of identical base pairs (liftover length x percent identity) of the total length of the original enhancer sequence. Bin size = 2%. **C-E.** Correlation analysis between the conservation of accessibility and the percent identity between the overlapping sequence only (C), overlap length (D) and the sequence homology relative to the length of the original sequence (E). Linear fit lines are shown in red, and the Pearson’s coefficient with its p-value are shown at the bottom right corner of each plot.

**Figure S2:**
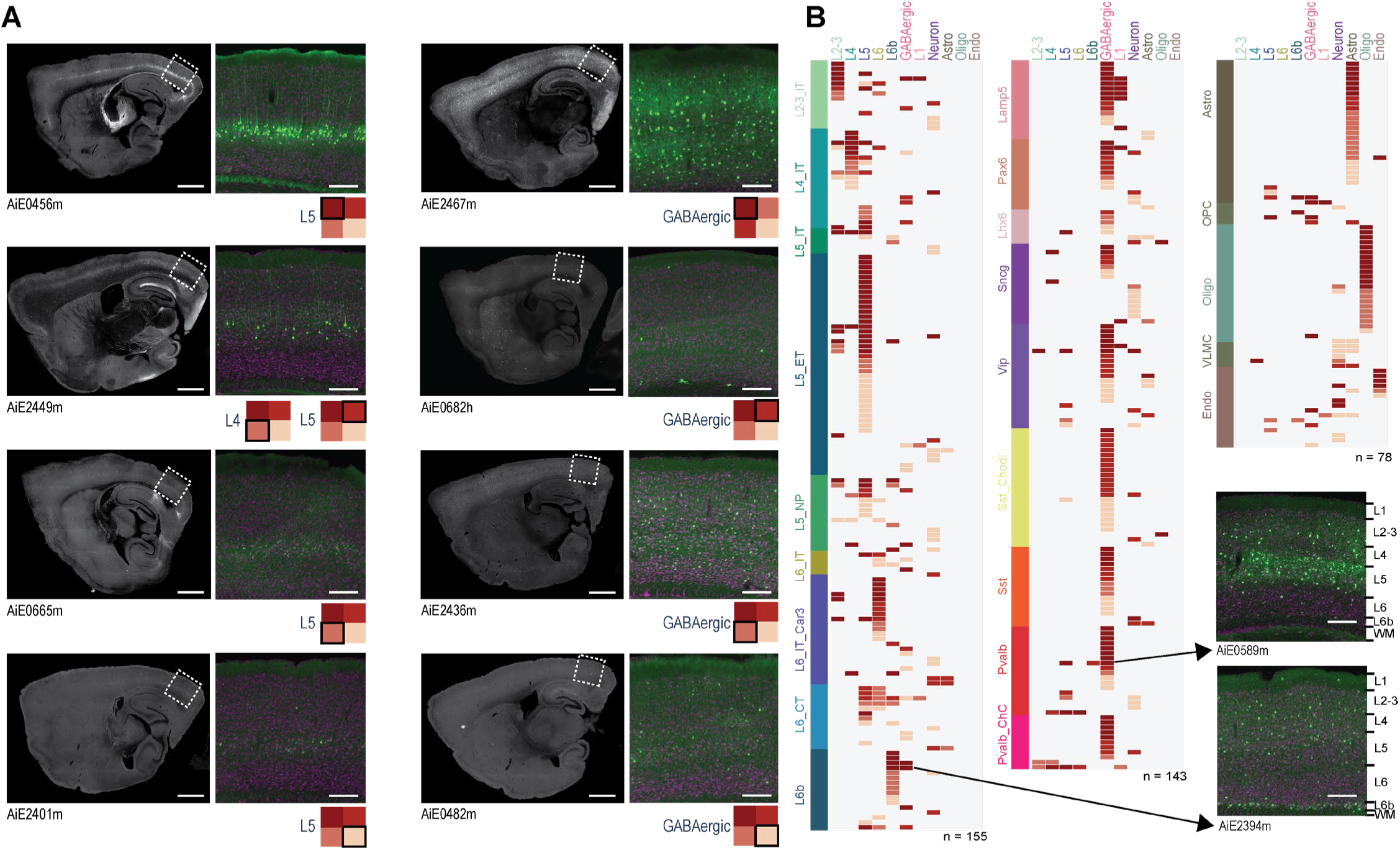
Enhancer scoring based on epifluorescence image sets. **A.** Representative sagittal images from eight experiments with focus on two cell categories: L5 (left column) or GABAergic (right). Scoring according to the scheme in Figure 3B is shown below each set of images. **B.** A heat map of all enhancers evaluated, which produced any labeling in the neocortex, arranged according to the cell population where their accessibility was highest, showing the identity of the labeled cell population and the labeling quality, according to the scheme in Figure 3B. The number of individual experiments in each category is shown below each plot (total n = 376). For images, scale bars for full section and expanded views are 1.0 and 0.2 mm, respectively.

**Figure S3:**
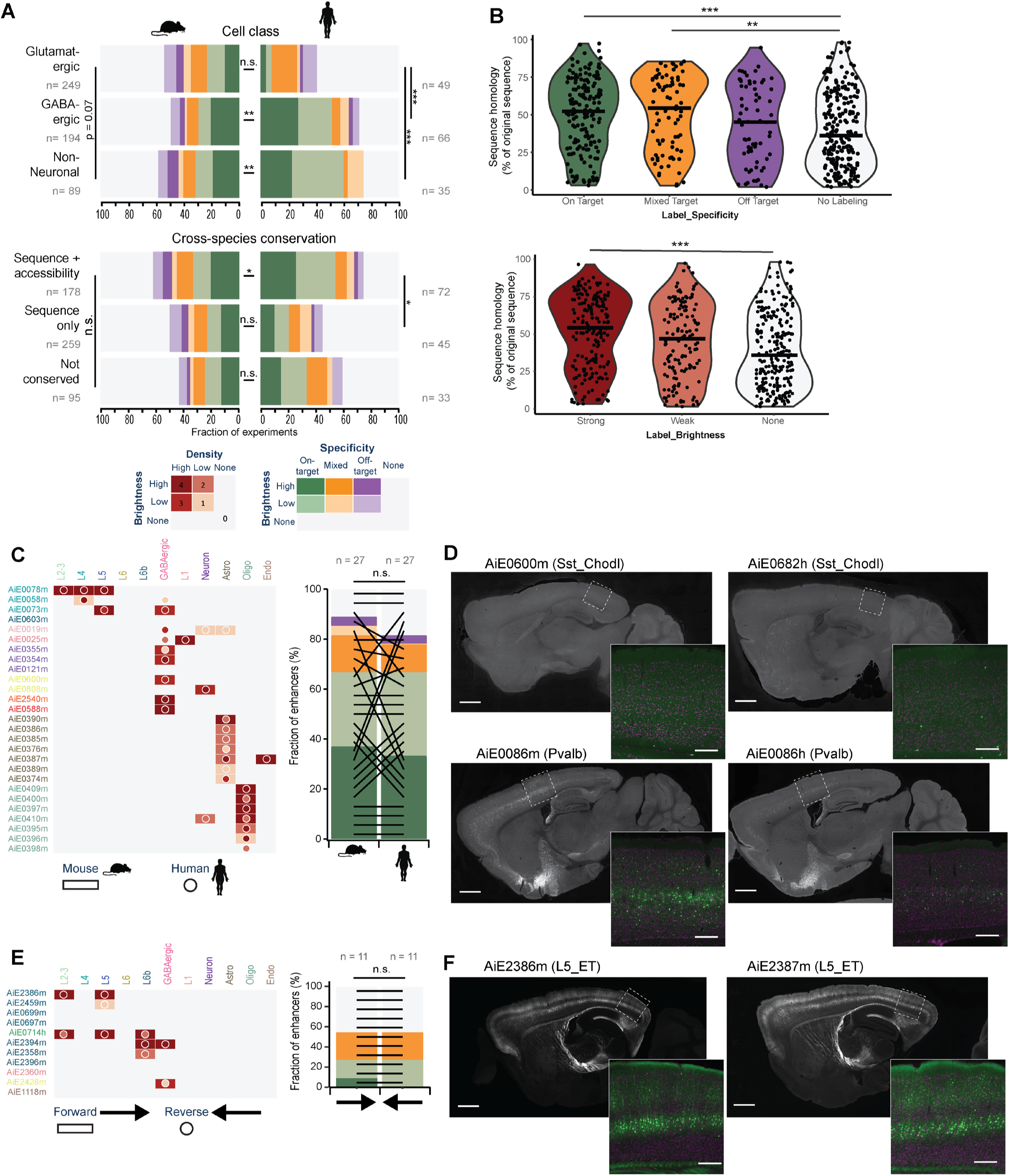
Effects of cross-species conservation and sequence orientation on enhancer performance. **A.** Summary plot of enhancer scoring data according to their genome of origin (top) and cross-species conservation of sequence and accessibility (bottom). **B.** Violin plots showing the degree of mouse-human sequence homology according to the different specificity (top) and brightness (bottom) categories. P-values were calculated with one-way ANOVA, followed by a pairwise post-hoc analysis (Tukey) corrected for multiple comparisons (Bonferroni). Significance levels are the same as in (A). **C.** Summary plot of the scoring data for mouse (rectangles) alongside its human (circles) orthologous sequence (Left) and a summary plot of the scoring data according to the brightness and specificity, with black lines connecting each pair (Right). **D.** Representative epifluorescence images of sagittal sections, showing labeling pattern for two individual mouse enhancers (left) and their human orthologs (right). **E.** Summary plot of the scoring data for complementary oriented sequences (left) and a summary plot of the scoring data according to the brightness and specificity, with black lines connecting each pair (right). **F.** Representative epifluorescence images of sagittal sections, showing identical labeling pattern for a complementary pair. P-values were calculated using a chi-squared test. P-values < 0.05, < 0.01 and < 0.001 are denoted by *, **, ***, respectively, n.s. = not significant. Scale bars for full section and higher-magnification view = 1.0 and 0.2 mm, respectively.

**Figure S4:**
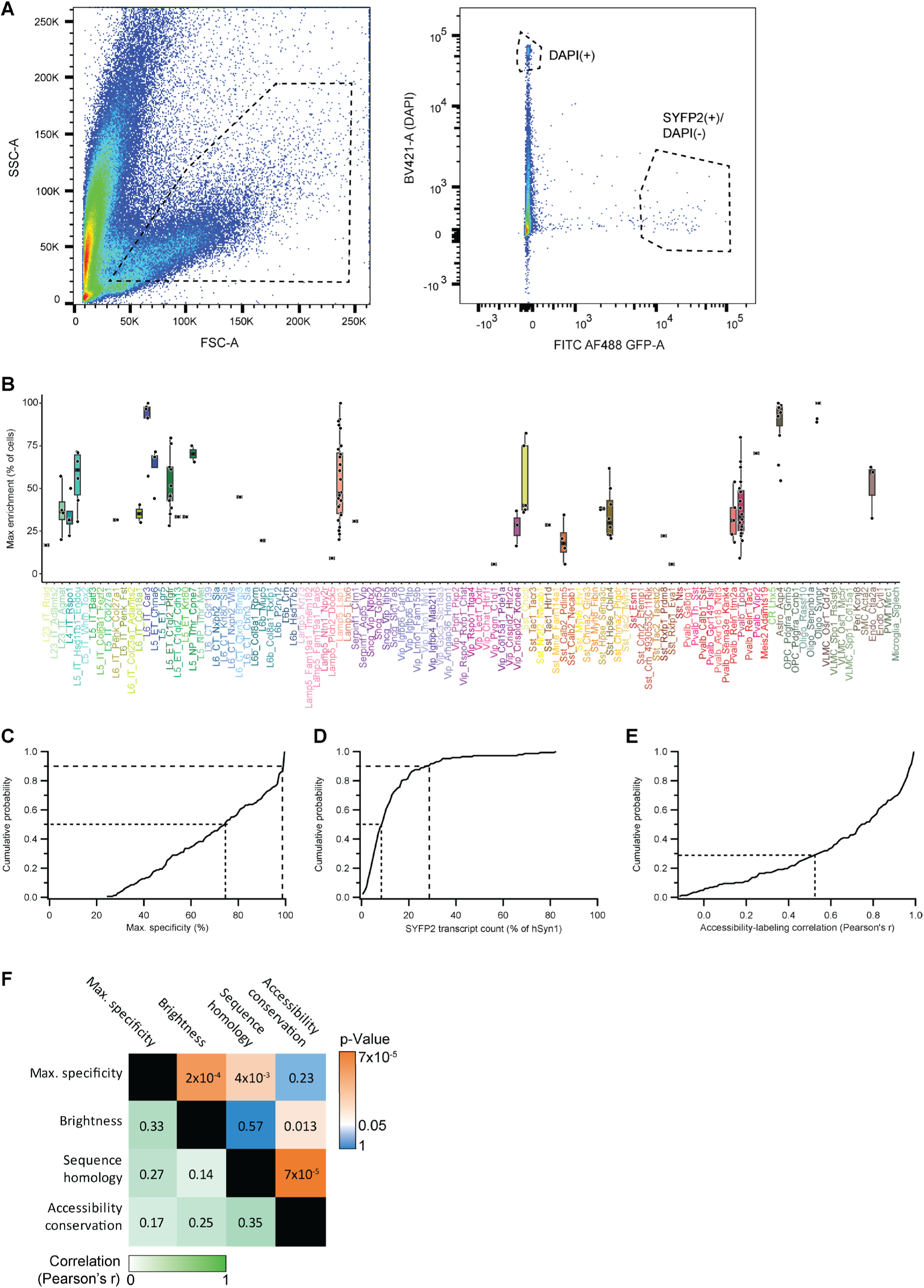
Distribution of specificity and brightness across enhancer AAVs. **A.** Representative plots of the FACS gating strategy for selective collection of cells labeled by the Lamp5 enhancer AiE2103m: Forward (FSC-A) and side scatter (SSC-A) were used to select objects matching size and granularity of cortical cells (left) and this fraction was further separated according to signal detected in the DAPI and FITC channels (right), in order to avoid collection of DAPI+ cells, whose membrane is likely compromised. Dashed boxes indicate the gates applied for sample collection. **B.** Box plots showing the distribution of enhancer maximum specificity in each of the cortical clusters. **C.** A cumulative distribution plot showing the fraction of enhancers as a function of their specificity, estimated by the maximal fraction of labeled cells. **D.** A cumulative distribution plot showing the fraction of enhancers as a function of their brightness, relative to hSyn1. **E.** A cumulative distribution plot showing the fraction of enhancers as a function of correlation coefficient, between the distribution of labeled cells and distribution of chromatin accessibility, across the cortical subclasses. **F.** Cross-correlation plot showing correlation values (white-green scale, bottom left corner) and their respective p-values (blue-orange scale, top right corner). Dashed lines in plots (A-C) show the median and top 10^th^ percentile of enhancers. P-values in (D) were corrected for multiple comparisons.

**Figure S5:**
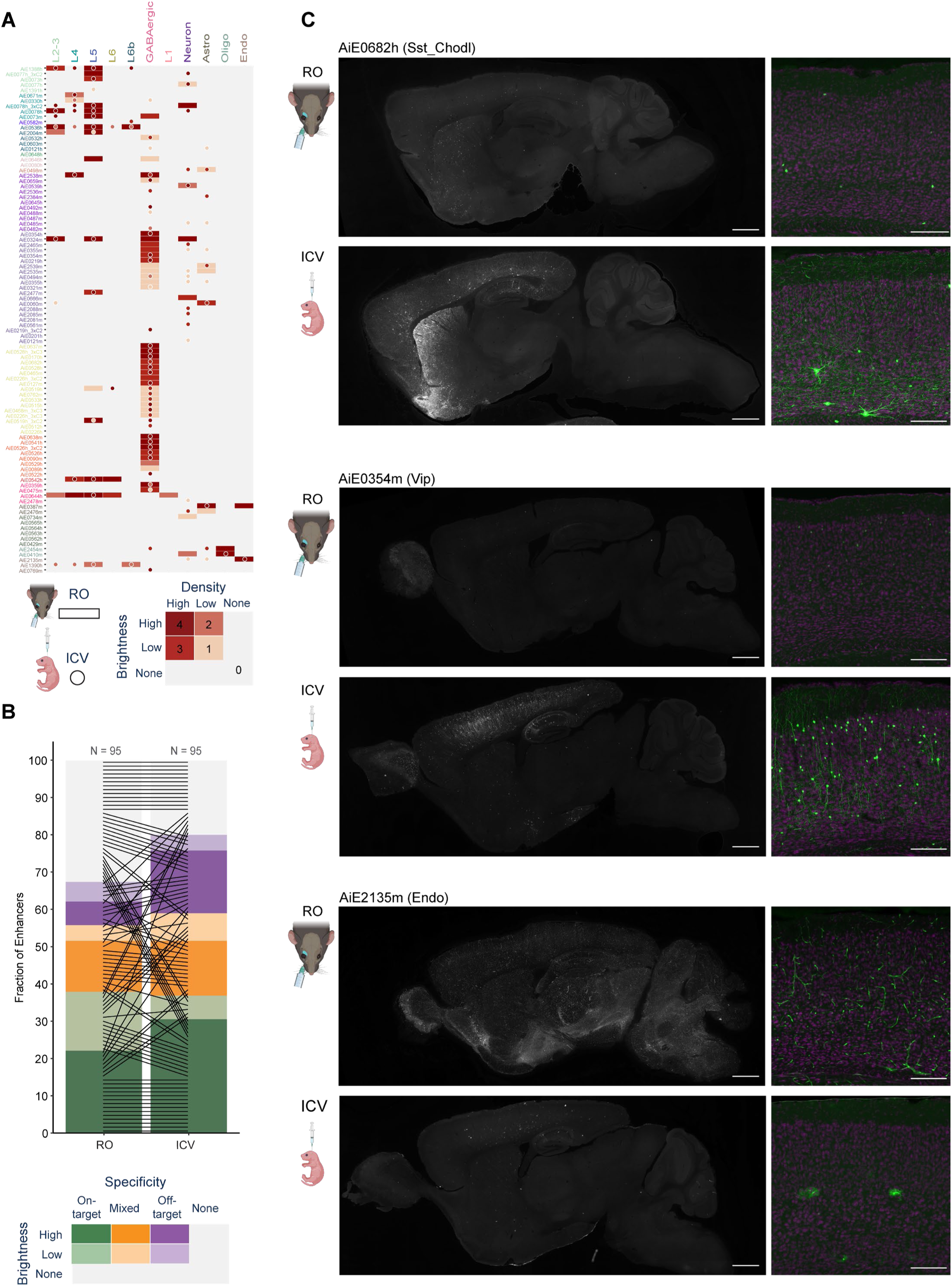
Comparison RO and ICV viral delivery routes by image scoring data. **A.** Heatmap of scoring data for the same vectors, delivered RO (rectangles) or ICV (circles). **B.** Summary plot of the scoring data according to the brightness and specificity, with black lines connecting each pair. **C.** Representative epifluorescence images of sagittal sections of three individual enhancers, comparing labeling pattern when the virus was delivered via the RO (top) or ICV (bottom) route. An expanded view of the visual cortex is displayed to the right of the full-sized image. Scale bars for full section and expanded view = 1.0 and 0.2 mm, respectively.

**Figure S6:**
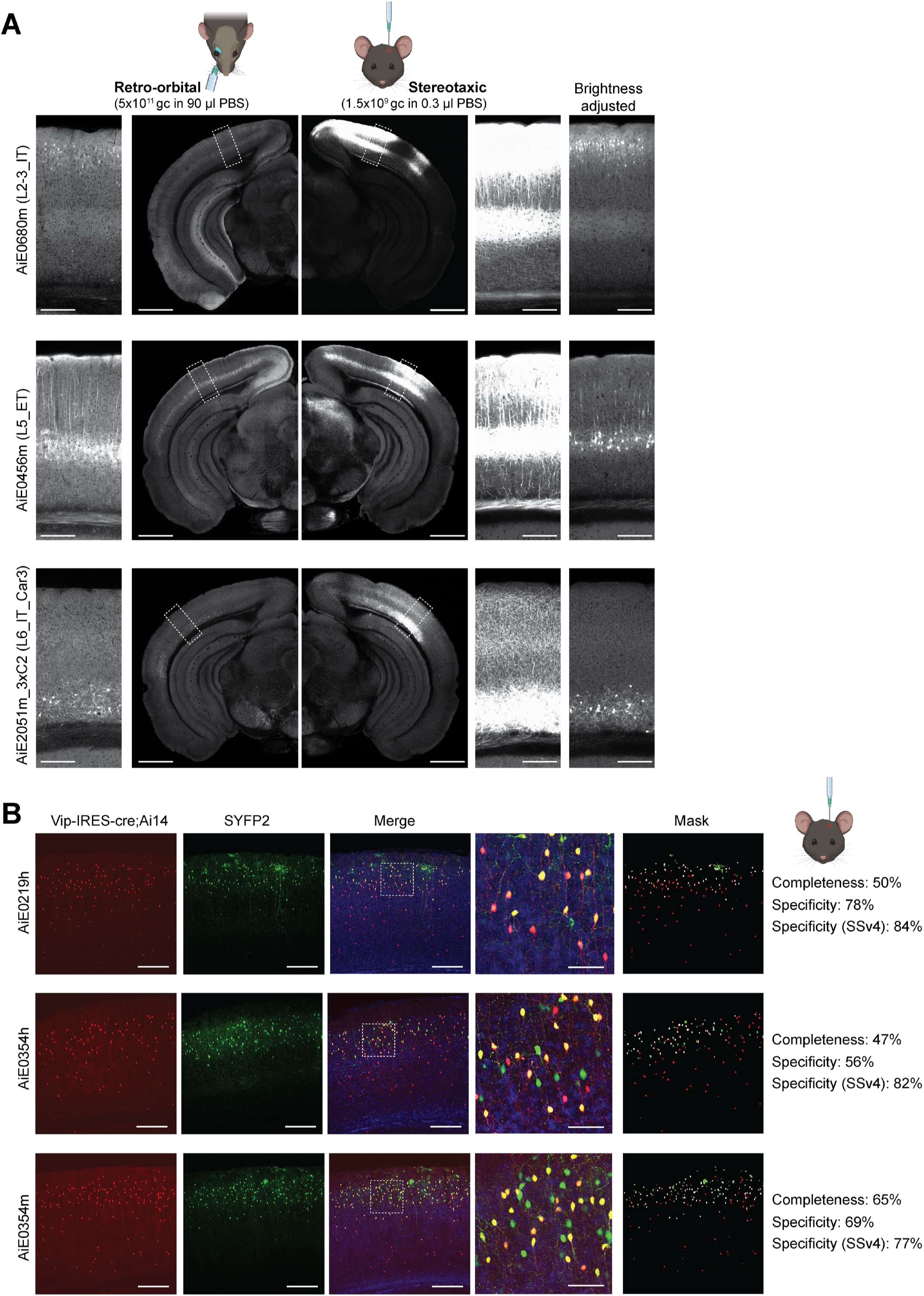
Stereotaxic delivery of enhancer AAVs. **A.** Stereotaxic delivery into VISp of three enhancers AAVs targeting different subclasses of glutamatergic neurons, resulted in strong, layer restricted SYFP2 expression. Scale bars for full section and expanded view = 1.0 and 0.2 mm, respectively. **B.** Stereotaxic delivery of three enhancers targeting Vip interneurons (green), delivered to the VISp of *Vip-IRES-Cre;Ai14* double transgenic line (red). For each injection, completeness was calculated as the fraction of SYFP2^+^/tdTomato^+^ cells, of all tdTomato^+^ cells at the injection site, and specificity was calculated as the fraction of SYFP2^+^/tdTomato^+^ cells, of all SYFP2^+^ cells. Specificity results were compared with SSv4 measurements for each vector, following RO delivery of 5×10^11^ genome copies (gc). n = 1 experimental animal for all experiments shown in this figure. Scale bars for full VISp view and expanded view = 0.2 and 0.05 mm, respectively.

**Figure S7:**
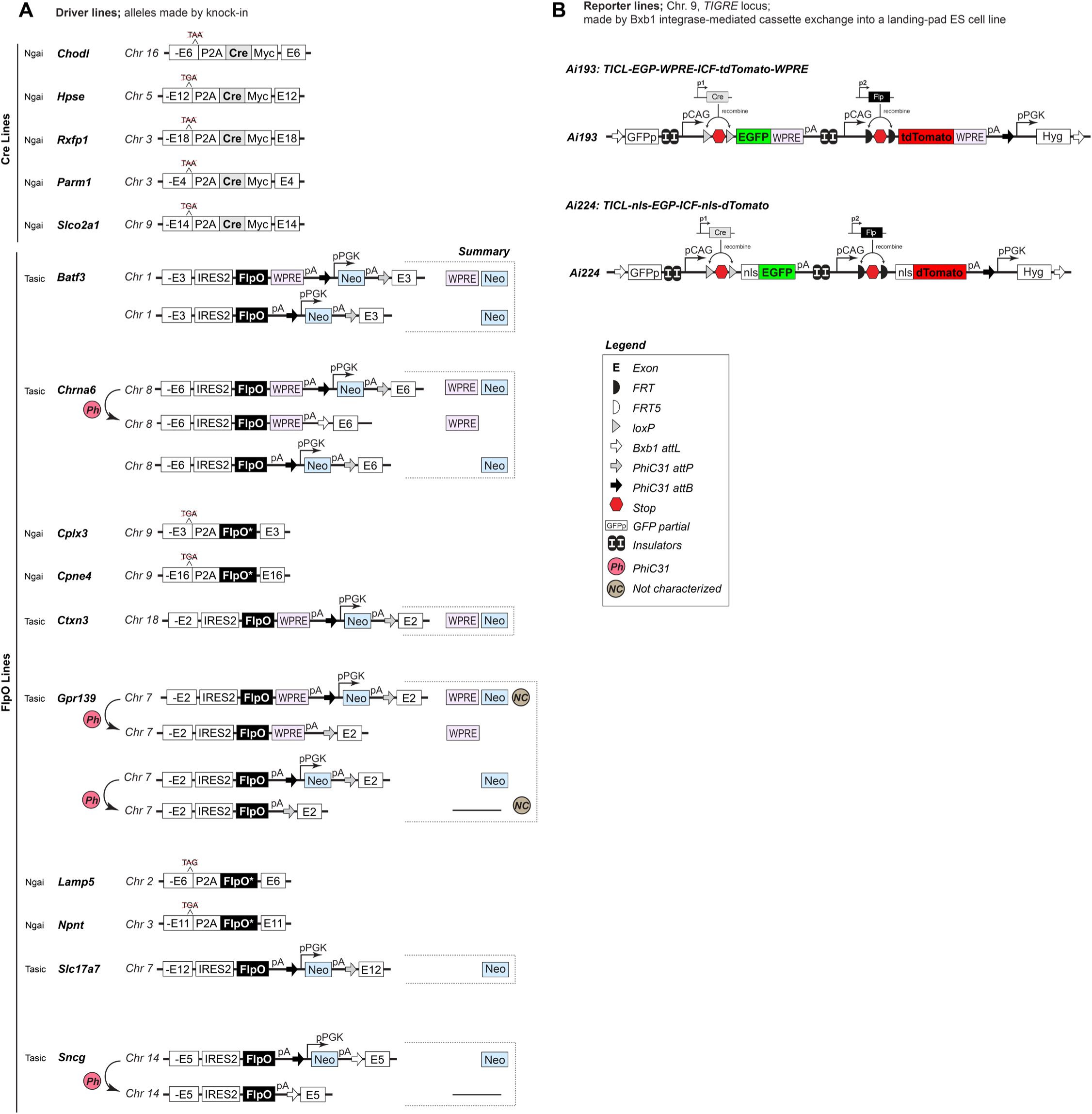
Transgenic line designs. **A.** Schematics depicting design of the 15 driver lines. Of these, five express Cre recombinase whereas 10 express FlpO. For some lines, such as *Chrna6-IRES2-FlpO*, we have versions with WPRE, with Neo present as well as with Neo removed allowing us to compare expression patterns in all three. In some instances, the driver lines were used as is and in others, they were crossed with *Rosa26-PhiC31* mice to delete the pPGK-neo selection cassette. **B.** Schematic depicting the design of the two new reporter mice *Ai193* and *Ai224*.

**Figure S8.**
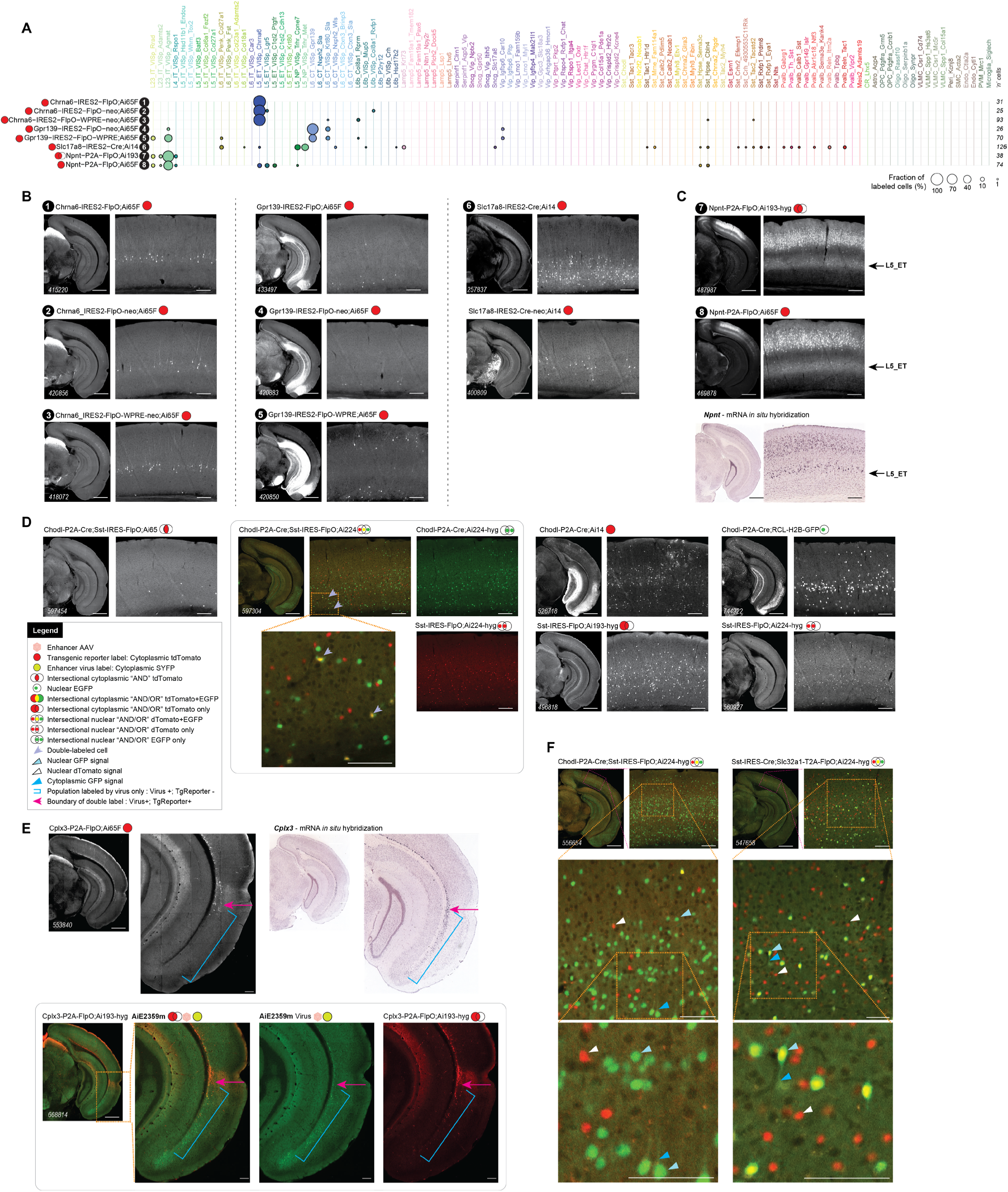
Factors influencing tool expression and evaluation. **A.** scRNA-seq (SSv4) data showing distribution of labeled cells from tools **1-8** mapped to mouse VISp taxonomy and displayed at the cluster level. **B.** Select STPT images for tools **1-6**, and additional related tools. **C.** Representative *Npnt* mRNA *in situ* hybridization and STPT images of *Npnt-P2A-FlpO* with two different reporters showing labeling of cells in L5, whereas the SSv4 data for the cross to *Ai193* (tool **7** in A) do not show L5 cells. This could be due to L5_PT cells not surviving FACS for this experiment. **D**. Representative STPT data for *Chodl-P2A-Cre; Sst-IRES-FlpO* crossed with previously characterized reporters (*Ai14* and *Ai65F*) and the new AND/OR reporters (*Ai193* and *Ai224*) both independently and as a triple transgenic. **E**. Representative STPT images showing *Cplx3-P2A-FlpO* with different reporters and *Cplx3* mRNA expression (blue brackets) by RNA *in situ* hybridization (https://mouse.brain-map.org/experiment/show/70928340). The expression pattern for the enhancer AAV, AiE2359m, mirrors *Cplx3* expression (blue brackets) by RNA *in situ* hybridization, whereas the expression of the transgenic line, *Cplx3-P2A-FlpO;Ai193* does not include *Cplx3+* cells in the entorhinal area. **F.** Expression of nls-EGFP (Cre-dependent) and nls-dTomato (Flp-dependent) is faithful in the *Ai224* reporter line; however, nuclear localization is imperfect. The GFP appears mostly nuclear, but weak signal can be observed in the cytoplasm (light blue arrow) and processes (blue arrow). In comparison, nls-dTomato appears nuclear (white arrow). Scale bars: 1.0 and 0.2 mm for full section and expanded view; 0.1 mm for further expanded view in (C).

## Supplemental tables

**Table S1.** Summary of labeling patterns produced by transgenic mouse lines, *related to Figures 7* and *8*.

**Table S2.** Transgenic mouse lines used in the manuscript, *related to STAR methods*.

**Table S3.** AAV plasmids used in the manuscript, *related to STAR methods*.

**Table S4.** Unique enhancers evaluated in this manuscript, *related to STAR methods*.

**Table S5.** “Hall-of-fame” enhancer AAVs, *related to Figures 3-6*.

